# Novel small molecules inhibit proteotoxicity and inflammation: Mechanistic and therapeutic implications for Alzheimer’s Disease, healthspan and lifespan- Aging as a consequence of glycolysis

**DOI:** 10.1101/2023.06.12.544352

**Authors:** Rachel Litke, James Vicari, Bik Tzu Huang, Lila Shapiro, Kun-Hyung Roh, Aaron Silver, Pooja Talreja, Nicolle Palacios, Yonejung Yoon, Christopher Kellner, Husnu Kaniskan, Sindhu Vangeti, Jian Jin, Irene Ramos-lopez, Charles Mobbs

## Abstract

Inflammation drives many age-related, especially neurological, diseases, and likely mediates age-related proteotoxicity. For example, dementia due to Alzheimer’s Disease (AD), cerebral vascular disease, many other neurodegenerative conditions is increasingly among the most devastating burdens on the American (and world) health system and threatens to bankrupt the American health system as the population ages unless effective treatments are developed. Dementia due to either AD or cerebral vascular disease, and plausibly many other neurodegenerative and even psychiatric conditions, is driven by increased age-related inflammation, which in turn appears to mediate Abeta and related proteotoxic processes. The functional significance of inflammation during aging is also supported by the fact that Humira, which is simply an antibody to the pro-inflammatory cytokine TNF-a, is the best-selling drug in the world by revenue. These observations led us to develop parallel high-throughput screens to discover small molecules which inhibit age-related Abeta proteotoxicity in a *C. elegans* model of AD AND LPS-induced microglial TNF-a. In the initial screen of 2560 compounds (Microsource Spectrum library) to delay Abeta proteotoxicity, the most protective compounds were, in order, phenylbutyrate, methicillin, and quetiapine, which belong to drug classes (HDAC inhibitors, beta lactam antibiotics, and tricyclic antipsychotics, respectably) already robustly implicated as promising to protect in neurodegenerative diseases, especially AD. RNAi and chemical screens indicated that the protective effects of HDAC inhibitors to reduce Abeta proteotoxicity are mediated by inhibition of HDAC2, also implicated in human AD, dependent on the HAT Creb binding protein (Cbp), which is also required for the protective effects of both dietary restriction and the *daf-2* mutation (inactivation of IGF-1 signaling) during aging. In addition to methicillin, several other beta lactam antibiotics also delayed Abeta proteotoxicity and reduced microglial TNF-a. In addition to quetiapine, several other tricyclic antipsychotic drugs also delayed age-related Abeta proteotoxicity and increased microglial TNF-a, leading to the synthesis of a novel congener, GM310, which delays Abeta as well as Huntingtin proteotoxicity, inhibits LPS-induced mouse and human microglial and monocyte TNF-a, is highly concentrated in brain after oral delivery with no apparent toxicity, increases lifespan, and produces molecular responses highly similar to those produced by dietary restriction, including induction of Cbp inhibition of inhibitors of Cbp, and genes promoting a shift away from glycolysis and toward metabolism of alternate (e.g., lipid) substrates. GM310, as well as FDA-approved tricyclic congeners, prevented functional impairments and associated increase in TNF-a in a mouse model of stroke. Robust reduction of glycolysis by GM310 was functionally corroborated by flux analysis, and the glycolytic inhibitor 2-DG inhibited microglial TNF-a and other markers of inflammation, delayed Abeta proteotoxicity, and increased lifespan. These results support the value of phenotypic screens to discover drugs to treat age-related, especially neurological and even psychiatric diseases, including AD and stroke, and to clarify novel mechanisms driving neurodegeneration (e.g., increased microglial glycolysis drives neuroinflammation and subsequent neurotoxicity) suggesting novel treatments (selective inhibitors of microglial glycolysis).

## Introduction

The dramatic increase in the use of antibiotics and vaccines during the 20^th^ century has led to a doubling of life expectancy from about 40 years in 1900 to about 80 years today, even excluding infant mortality without, unfortunately, a concomitant increase in healthspan [1]. Today morbidity and mortality are largely due to age-related chronic diseases, including dementias caused by Alzheimer’s, Parkinson’s, Huntington’s, and vascular Diseases, which generally entail, and are plausibly driven by, both age-related increased microglia inflammation [2] [3] and age-related proteotoxicity [4, 5], the former likely mediating the latter. Furthermore, dietary restriction (DR), which generally delays age-related diseases and increases lifespan, inhibits inflammation (innate immunity) in humans and other species [6], and evidence supports that inhibition of inflammation mediates protective effects of DR [46] [7]. The functional significance of inflammation during aging is also supported by the fact that Humira, which is simply an antibody to the pro-inflammatory cytokine TNFa, is the best-selling drug in the world by revenue, even though it does not cross the blood-brain barrier [8]. Furthermore, both inflammation [9] [10] and proteotoxicity [11], like cancer [12], entail and are arguably driven by increased glycolysis (the Warburg effect), and considerable evidence suggests that protective effects of DR are mediated by reduced glycolysis, reversing the Warburg effect [13].

Dementia, especially due to Alzheimer’s Disease (AD), arguably produces the greatest strain of any condition on world-wide health systems, up to $1 trillion in 2018, affecting more than 50 million people [14]. The emotional toll on caregivers is also substantial [15] and there is generally agreement that dementia substantially reduces quality of life of both patients and caregivers [16]. Until recently four individual drugs and one combination drug have been approved for use in patients with AD [17], but the effect sizes of these drugs are relatively modest and the clinical significance of these effects, especially whether the benefits outweigh the harm, have been widely questioned [18]. For example, Eliti [19] simply states that “the efficacy of these is widely debated.” The limited efficacy of the approved drugs is probably because these treatments address symptomatic relief, rather than addressing causes of dementia. Clinical trials of other therapies, primarily targeting Abeta, have generally failed [20] [21]. More recently the FDA approved an antibody against Abeta aggregates, but this was a highly controversial approval and scientific consensus is that this antibody is not clinically effective [22, 23].**Thus development of pharmaceutical treatments for dementia which produce clinically significant amelioration of the condition, addressing the causes of dementia, is of the utmost priority.**

AD is clinically defined according to post-mortem analysis which quantifies the density of silver-stained plaques (consisting primarily of aggregates of the Abeta peptide) and tangles (consisting primarily of hyperphosphorylated Tau). There is general consensus that proteotoxicity involving both Abeta and Tau and possibly other proteins is at least partly responsible for driving AD pathophysiology [24]. Since neurodegeneration is the hallmark feature of AD, the neuron has understandably been the focus of AD research since the discovery of the disease by Alois Alzheimer [25]. Nevertheless, recent genetic studies have clearly implicated microglial activity, which serve as the primary driver of innate immunity in the brain, as a key driver of the disease [26]. This activity is plausibly at least somewhat independent of Abeta [27], although Abeta1-42 peptide does produce microglial release of TNFa and IL-6 [28]. Of particular significance, numerous studies have demonstrated that inhibiting TNFa ameliorates impairments in rodent models of AD [29] [30, 31] and even, according to case reports, in humans [32] [27, 29-50]. Particularly compelling, chemical ablation of microglia improves outcome in the 3XTg mouse model of AD [51], strongly implying that microglial hyperactivity (likely including increased secretion of TNF-α) plays a major role in driving the pathophysiology of AD.

These observations led us to develop parallel high-throughput screens to discover small molecules which inhibit proteotoxicity, inflammation, and glycolysis, increase lifespan, which cross the blood-brain barrier after oral delivery with minimum toxicity, is highly protective in a mouse model of stroke. The effects of these small molecules were used to further develop evidence on mechanisms driving the pathophysiology of dementia with implications for further development of small molecules protecting against dementia.

## Results

### Initial screen for compounds which delay Abeta proteotoxicity in vivo

Our initial screen used the Microsource Spectrum Collection (2560 compounds) in a *C. elegans* model (CL2006) of transgenic human Abeta1-42 proteotoxicity, which we [52] and others [53] have used to probe mechanisms mediating protective effects of DR on proteotoxicity in the context of the whole animal, since such effects may be cell non-autonomous [5]. This screen entailed administering each compound at 8 uM (lower concentration than in similar C. elegans screens [54]) in liquid medium cultivated with *C. elegans* strain CL2006 beginning at Adult Day 1. Abeta-induced proteotoxicity was quantified as paralysis at Adult Days 10, 11, 12, at which time about 97% of the CL2006 strain are paralyzed (wild-type *C. elegans* never exhibit this phenotype). Remarkably, the most protective compound (exhibiting only 12% paralysis compared to 97% controls) was phenylbutyrate (PBA), an HDAC inhibitor approved as a therapy for urea cycling disorder, an orphan disease, and which is being developed as a possible therapy for a wide range of neurological conditions [55] including AD [56-62]. An early indication of the potential for HDAC inhibitors to treat AD was our previous report that a structurally related HDAC inhibitor, sodium butyrate, mimicked the protective effects of DR to delay impairments in the CL2006 *C. elegans* model of Abeta proteotoxicity, and increase lifespan in wild-type *C. elegans* [52]. These observations have been corroborated and extended [63], including the demonstration that Cbp expression in GABAergic neurons double the protective effects of Cbp to increase lifespan are mediate, We examined the effect of butyrate on lifespan because we had demonstrated that induction of the histone acetyltransferase Creb-binding protein (Cbp) is essential for the protective effects of dietary restriction on lifespan and delay of Abeta proteotoxicity, and indeed inhibition of Cbp also blocked the protective effects of the HDAC inhibitor butyrate [52]. Since butyrate and phenylbutyrate inhibit several HDACs, we carried out an RNAi screen to determine which HDAC was most implicated in driving impairments in the C. elegans model of Alzheimer’s Disease. These studies indicated that RNA I inhibition of HDAC2 was most protective in delaying impairments in this model (**Fig. 1A**). We then screened several small molecules that were relatively specific to specific HDACs; the most protective of these HDAC inhibitors was Kevetrin, a relatively specific inhibitor of HDAC2 and HDAC6 (**Fig. 1B**). This result was of particular interest because brain HDAC2 is preferentially induced with age, an effect blocked by dietary restriction [64], and is preferentially induced in patients with AD and in animal models of AD [65]. Similarly genetic ablation of HDAC1 and HDAC2 specifically in adult microglia delays impairments in the 5XFAD mouse model of Alzheimer’s Disease (Datta et al., 2018). Furthermore, genetic ablation of HDAC2 in microglia, either genetically or pharmacologically, promotes the transition from the proinflammatory state to the M2 phagocyotic state and is highly protective in a mouse model of stroke (Yang et al, 2021).

**Figure 1.**
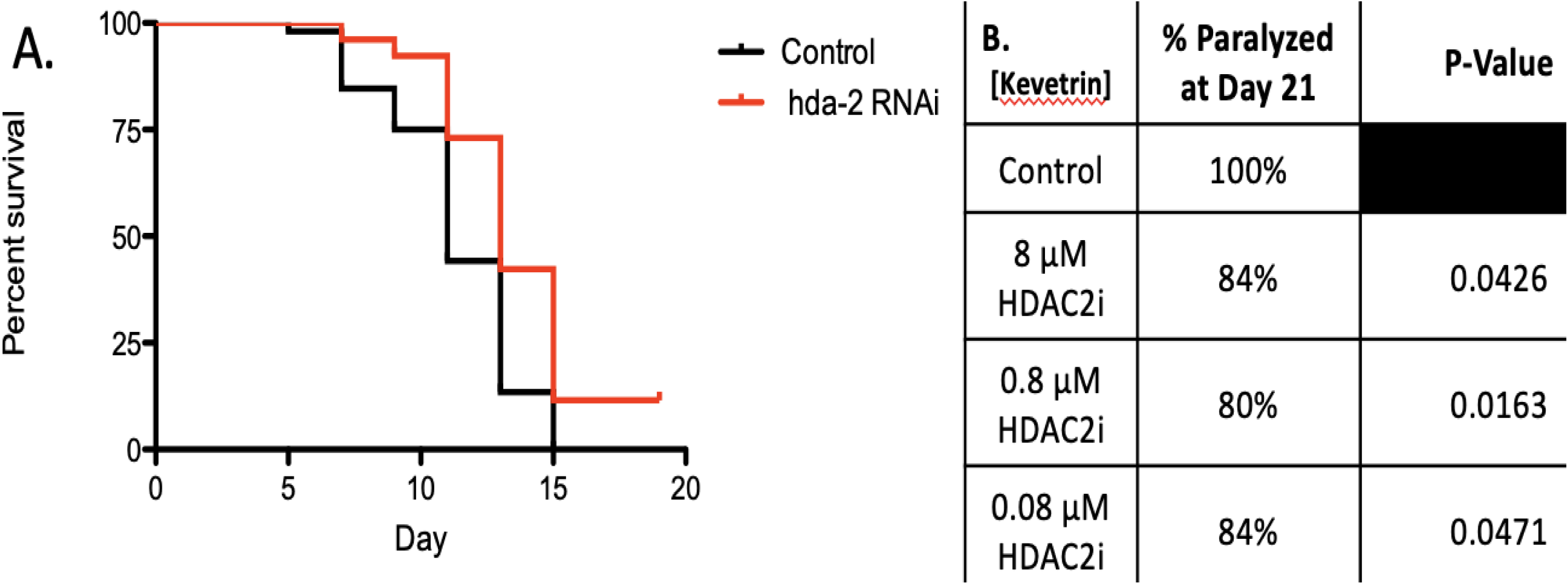
**(A) RNAi inhibition of HDAC2, but not RNAi of other HDAC isoforms, significantly (p<0.001) delayed impairments in a C. elegans model of human Abeta 1-42 (CL2006). (B) Kevetrin, a relatively specific HDAC2 inhibitor, but not other inhibitors relatively specific to other HDAC isoform, significantly (p-val es shown in Table) across a range of concentrations.**

The second-most protective drug to delay Abeta proteotoxicity in the C. elegans CL2006 model was the beta lactam antibiotic methicillin (17% paralyzed vs. 97% paralyzed controls). Beta lactam antibiotics are protective in a wide range of neurological conditions in animal models, including in animal models of AD [66-69] [66-68], although the mechanisms mediating these protective effects are not fully elucidated. We therefore assessed if other beta lactam antibiotics would also delay Abeta protoxicity in the C. elegans CL2006 model. These studies indicated that several other beta lactam antibiotics, including Cefproxil, Ampicillin, and Amoxicillin, also delayed Abeta proteotoxicity (**Fig. 2**) as well as microglial secretion of TNFa. These results were consistent with with previous reports that beta lactam antibiotics significantly inhibited microglial inflammation (Zhou et al., 2021) independent of their antibiotic activity (Lim et al., 2021).

**Figure 2.**
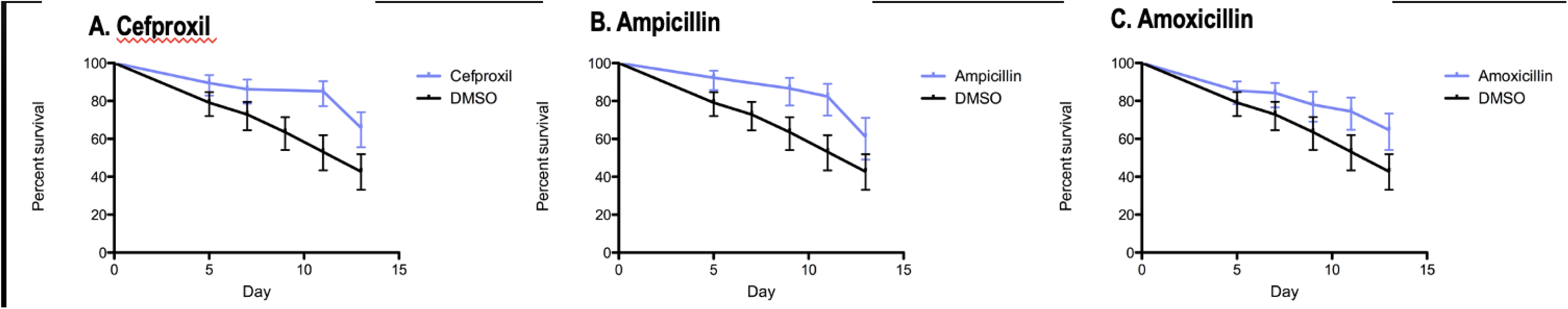
**Beta lactam antibiotics significantly (p<0.001) delay Abeta proteotoxicity (% non-paralyzed/% survival) in C. elegans CL2006 model.**

Finally, the third-most protective drug was quetiapine, a tricyclic atypical antipsychotic sometimes also used to treat depression (18% paralyzed vs. 97% paralyzed controls). As with the first 2 protective drugs, quetiapine delayed cognitive impairments and several indicators of pathology in a mouse model of Alzheimer’s Disease [70]. Most remarkably, after these analyses were completed and IP filed, a subsequent independent and novel bioinformatic analysis concluded that “antipsychotic drugs with circadian effects, such as Quetiapine … will have efficacious effects in AD patients” [71]. This observation is consistent with the high prevalence of both psychosis [72], which actually predicts the progression of cognitive impairments [73], and depression [74] [75-78] in patients with Alzheimer’s Disease. However, in clinical trials atypical antipsychotic drugs actually *accelerated* cognitive impairments [79]. This observation would appear to rule out treating AD with currently available atypical antipsychotic drugs. However, the results in mice suggested that compounds with similar bioactivities, but possibly fewer or at least different iatrogenic effects, might still prove to be effective in treating the disease. Furthermore, the high degree of co-morbidity of neuropsychiatric diseases and AD raise the interesting possibility these conditions may share some common pathophysiological mechanisms.

One such mechanism could plausibly involve neuroinflammation driven by microglial activation. GWAS have convincingly demonstrated that microglia activity is a major factor in the pathophysiology of AD [27] [46]. Microglia are the resident myeloid cells in the brain, and like myeloid cells (macrophages) in the periphery, when activated they undergo a process entailing inflammation (the M1 phase entailing secretion of cytokines and reactive oxygen species) followed by repair (the M2 phase entailing phagocytosis). Many lines of evidence strongly implicate microglial inflammation in the pathophysiology of AD [31, 33, 42, 45, 80-83]. Of particular interest, several reports have concluded that inhibitors of TNF-alpha would plausibly be protective in AD [29, 30, 32, 38–40, 84, 85]. However, these inhibitors are generally biologicals (antibodies and protein binders of TNF-alpha) so have limited capacity to cross the blood-brain barrier, suggesting that small molecule inhibitors of TNF-alpha secretion could be far more effective. Almost as extensive are the many studies implicating IL-6 in driving pathophysiology of AD [31, 33, 37, 41, 43, 44], although evidence that inhibiti g IL-6, in contrast to TNF-alpha, would ameliorate impairments in AD is lacking. Of course proteotoxicity, particularly driven by Abeta and Tau, also contribute to the pathophysiology of AD, but a strong case can be made that this proteotoxicity is mediated at least in part by neuroinflammation [49]. It is therefore of great interest that pro-inflammatory cytokines, especially IL-6 and TNFa, have been increasingly implicated in the development of neuropsychiatric conditions [86], including depression [87] and psychosis [88, 89].

Tricyclic compounds constitute the major class of drugs to treat psychosis and depression [90]. The combination of two tricylic antipsychotic phenothiazines, chlorpromazine and promethazine, is reported to be neuroprotective in a rat model of stroke associated with reduction in neuroinflammation [91, 92]. It is of particular interest that this reduction in neuroinflammation appears to be mediated by a reduction in “hyperglycolysis” [93] since reduction of glycolysis appears to play a major role in mediating the protective effects of dietary restriction during aging [13, 94-96]. Furthermore, increased glycolysis appears to be a requirement for myeloid cells to support inflammation (likely due to the requirement to produce reactive oxygen species) [97-99], just as increased glycolysis appears to be a requirement for the development of cancer (the Warburg effect).

Our initial screen involved screening the Microsource Spectrum Collection (2560 compounds) to discover compounds which would delay impairments in a C. elegans model (CL2006) of proteotoxicity in Alzheimer’s Disease due to transgenic expression of the human Abeta 1-42 gene [53]. We have used this model to probe mechanisms mediating protective effects of DR on proteotoxicity in the context of the whole animal, since such effects may be cell non-autonomous [5]. This screen entailed maintaining the *C. elegans* beginning at Adult Day 1 in 384-well plates with liquid media containing 10 uM of each compound or vehicle at an E. coli concentration of 0.8 × 10^9^/ml, then filming at Adult Days 10, 11, and 12 using a 3-D printer based microscopic system designed for this purpose. Under these circumstances the vehicle controls were 100% paralyzed, so analysis for paralysis focused on Adult Day 12.The most protective compound (exhibiting only 12% paralysis compared to 97% controls) was phenylbutyrate (PBA), an HDAC inhibitor approved as a therapy for urea cycling disorder, an orphan disease, and which is being developed as a possible therapy for a wide range of neurological conditions [55]. The potential for HDAC inhibitors to treat AD had been suggested by our previous report that a structurally related HDAC inhibitor, sodium butyrate, mimicked the protective effects of dietary restriction to delay impairments in the CL2006 *C. elegans* model of Abeta model proteotoxicity, and to increase lifespan in wild-type *C. elegans* [52]. This report was followed by reports that PBA actually *reverses* cognitive impairments in mouse models of AD [56-62]. Also, as in *C. elegans*, PBA increases lifespan in Drosophila [100]. PBA is now in clinical trials with encouraging results on markers of AD (Amylyx). The second-most protective compound was methicillin (17% paralyzed), consistent with the many studies demonstrating that beta lactam antibiotics show great promise in treating AD and other neurological conditions, including stroke [66-69]. The third most protective compound was quetiapine (18% paralyzed), an atypical antipsychotic drug which improves symptoms in mouse models of AD and other forms of dementia [70, 101, 102], including patients with AD. These 3 results alone greatly support that this screening protocol in *C. elegans* can at least in principle lead to the development of small molecule therapies which could plausibly be useful to treat human dementias.

As described above, microglial gene expression and in particular microglial secretion of inflammatory cytokines such as TNFa, are increasingly implicated in driving the pathophysiology of AD [29-40, 84, 85, 103] and stroke [104-107]. A different cytokine, IL-6, has also been implicated in driving the neuropathology of both AD and stroke [108], suggesting that dementia in both conditions might be driven by a general increase in inflammatory cytokines. NFkappaB, the key transcription factor which generally induces pro-inflammatory cytokines, has also been directly implicated in the pathophysiology of AD [109] and stroke [110].

We therefore re-screened the 40 compounds which produced the most robust inhibition of Abeta-induced proteotoxicity in the *C. elegans* studies described above (from 12% to 33%, paralyzed, compared to 100% paralyzed in vehicle-treated controls) for efficacy to inhibit BV2 microglial secretion of TNFa. BV2 cells were exposed to 5ug/ml of LPS for 24 hours, followed by exposure to 8 uM of the test compound which in our first screen inhibited Abeta-induced proteotoxicity. After supernatant was removed to measure cytokine release, cell viability was assessed using the MTT assay [111]. in our hands stimulating inflammation by adding LPS (1.25-5ug/mL) was a more reliable stimulus of inflammation than hAbeta1-42 [99].

As shown in **Figure 3**, 23 of the 40 compounds which inhibited Abeta toxicity in the *C. elegans* model of AD also inhibited TNFa secretion (normalized to MTT). This far from random overlap greatly supports that inhibition of the innate immune system (inflammation) mediates the protective effects of these compounds to reduce proteotoxicity. Further supporting this hypothesis, the two most inhibitory compounds (#3 and #77), and the fifth most inhibitory compound (#4) were cortisol, flumethasone, and dexamethasone, classic glucocorticoids which are best known for, and whose main clinical use is for, their robust inhibition of innate immunity. It is perhaps surprising that these compounds were also highly effective to inhibit proteotoxicity in *C. elegans*, but it is well-established that these compounds exert protective effects in this species, at least in part by activating the longevity-enhancing gene Daf-16 [112], which turn is known to inhibits proteotoxicity in a *C. elegans* model of AD [113]. The other two most inhibitory compounds, #61 (Berbamine) and #18 (Emodin) have already been demonstrated to protect in mouse models of AD [114] [115] further validating the predictive power of our parallel phenotypic screening protocol.

**Figure 3.**
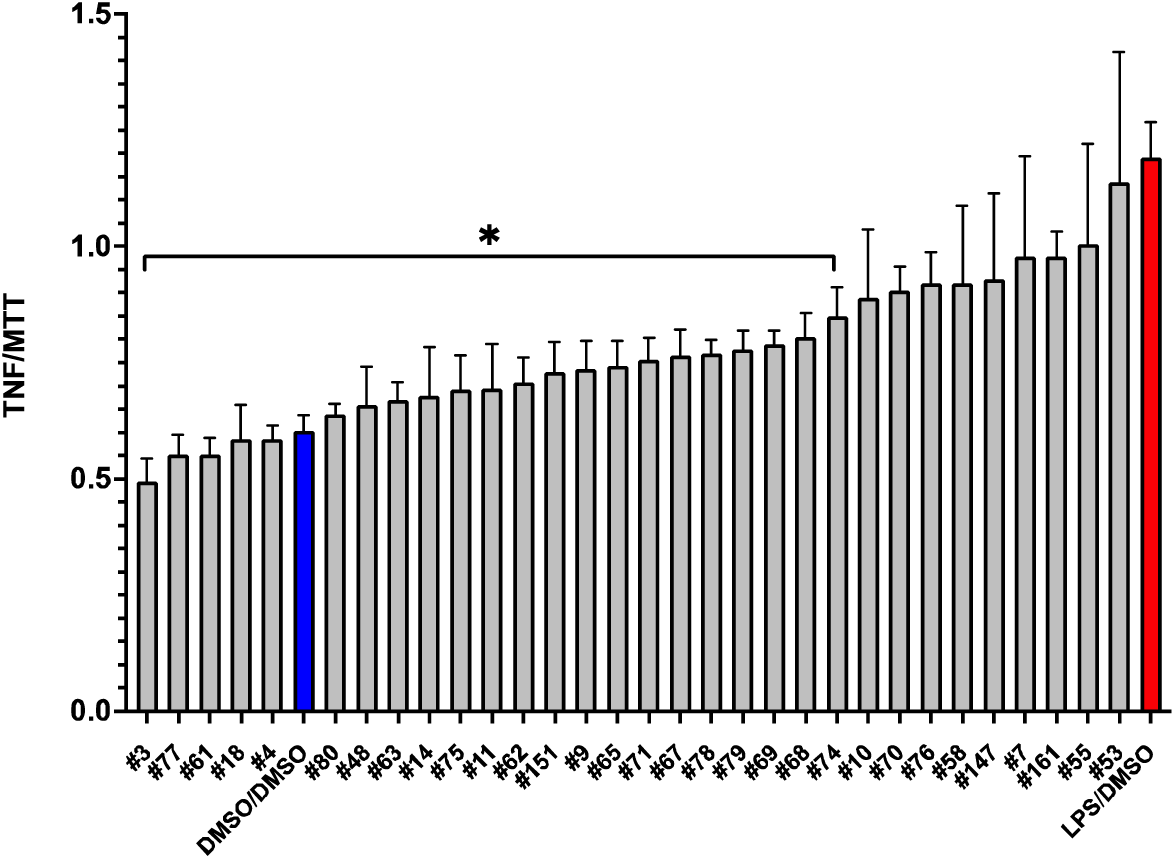
Inhibition of secreted TNFa in LPS-stimulated microglia treated with most protective compounds. Secreted TNFa was evaluated in BV2 microglia stimulated with 5ug/mL LPS for 24 hours followed by 8uM drug for 24 hours using ELISA (A.). Selected compounds were the top protective compounds from pretreatment studies. \**P* < 0.05 vs LPS condition. **(Only compounds which were not toxic were included in the analysis).**

The other compounds which inhibit Abeta proteotoxiciy and TNFa secretion represent several classes of drugs approved to treat a wide range of different health conditions. Two of the 4 most inhibitory compounds were the tricyclic compounds chlorprothixene (#45) and triflupromazine (#75), which like quetiapine are approved to treat psychosis. Chlorprothixene is a second-generation antipsychotic drug which in at least one study was the second-most prescribed drug for psychosis [116], and also effective in treating depression [117]. Triflupromazine is also approved to treat psychosis [118], and of particular interest also inhibits inflammation by blocking the association between IKKBeta and NEMO (NFkappaB Essential Modulator), thereby blocking the activation of the key pro-inflammatory transcription factor NFKappB) [119]. The ability of triflupromazine to block this activation of NFKappB was predicted based on structural features of this proinflammatory interaction and consistent with the critical role of inflammation driving dementia, NFKappaB is also widely implicated in driving AD and other dementias [120]. <u>We therefore propose that the most likely direct target of all the tricyclic compounds which</u> <u>block microglial cytokine secretion, including GM310 (see below) similarly do so by blocking this transcriptional</u> <u>complex.</u> In any case the remaining non-steroid, non-tricyclic compounds constitute a diverse class of lead compounds with also extremely promising properties to treat dementia. <u>Therefore the proposed studies</u> <u>based on these lead compounds would follow the same highly successful protocols we used to</u> <u>develop novel tricyclic compounds to treat dementia, to</u> <u>develop even more potent and safer novel compounds in other</u> <u>chemical classes to treat dementia.</u>

A key property of all potential treatments for dementia is the requirement to cross the blood-brain barrier after oral administration to be effective in psychiatric conditions, they are plausible candidates to treat dementia, offering a much more effective (and certainly less expensive) alternative to currently approved anti-inflammatory treatments such as Humira, the best-selling drug in the world by revenue. If so structure-activity relationship of congeners of these tricyclic compounds could lead to the development of novel compounds which cross the blood-brain barrier after oral delivery and could broadly reduce inflammation and proteotoxicity, and even potentially elucidate mechanisms driving age-related neurodegenerative as well cognitive impairments and neuropsychological conditions. We therefore assessed if congeners to these tricyclic compounds from the NCI compound library [121] would reduce microglial inflammation (indicated by reduced TNFa secretion as described above). Remarkably, all of these congeners also reduced TNFa secretion from BV2 microglia without producing toxicity (as indicated by the MTT assay), although the results were significant for only about half of these compounds (**Figure 4**).

**Figure 4.**
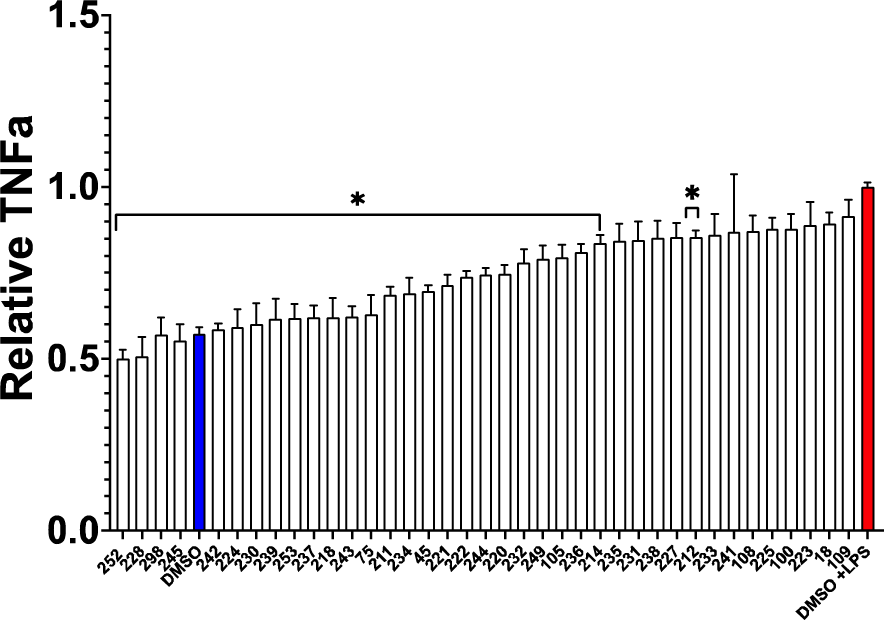
Inhibition of secreted TNFa in LPS stimulated microglia treated with tricyclic congeners of quetiapine from the NCI library of compounds. Secreted TNFa was evaluated in BV2 microglia stimulated with 5ug/mL LPS for 24 hours followed by 8uM drug for 24 hours using ELISA (A.). None of the drugs affected viability as measured by MTT.

To further probe protective effects of tricyclic antipsychotic compounds, we assessed efficacy of such compounds to delay Abeta proteotoxicity in the *C. elegans* CL2006 model. Remarkably, every tricyclic assessed in this assay significantly delayed Abeta proteotoxicity in this *C. elegans* model of AD (**Figure 5**), confirming and extending the observation of the protective effect of quetiapine in the same model.

**Figure 5.**
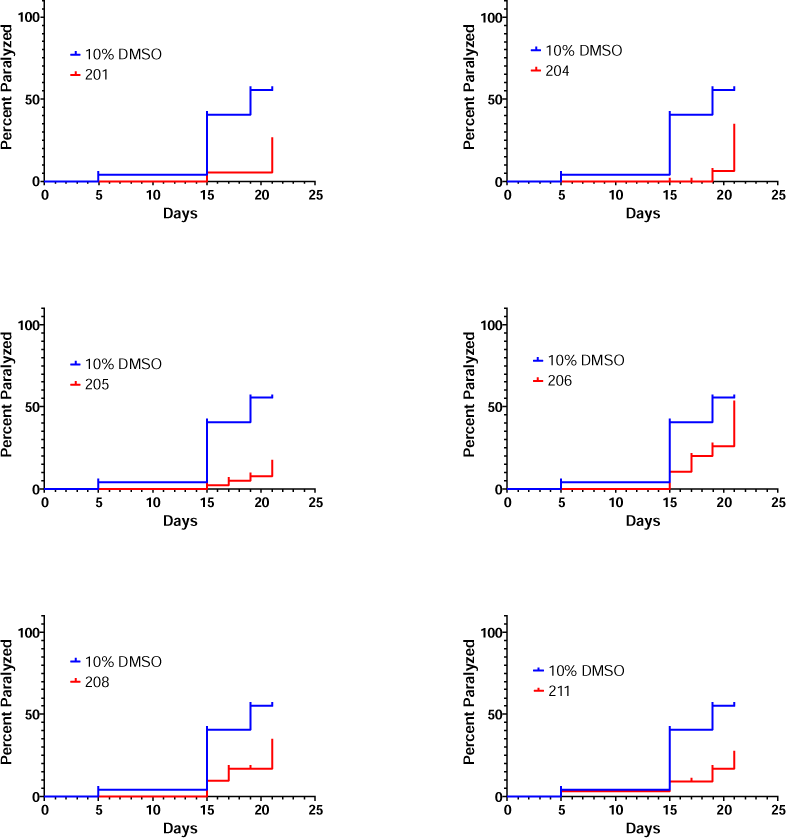
FDA-approved tricyclic antipsychotic drugs delay proteotoxicity in a *C. elegans* model of AD. All results are statistically significant (p<0.05).

To develop novel compounds even more protective and without detectable toxicity, we used the results from these structure-activity relationships to predict novel tricyclics predicted to exhibit such protective properties, which were then synthesized and assessed for their protective properties. As shown in **Figure 6**, novel tricyclic compound GM310 produced the greatest inhibition of TNF-a secretion without producing toxicity, as indicated by the MTT assay.

**Figure 6.**
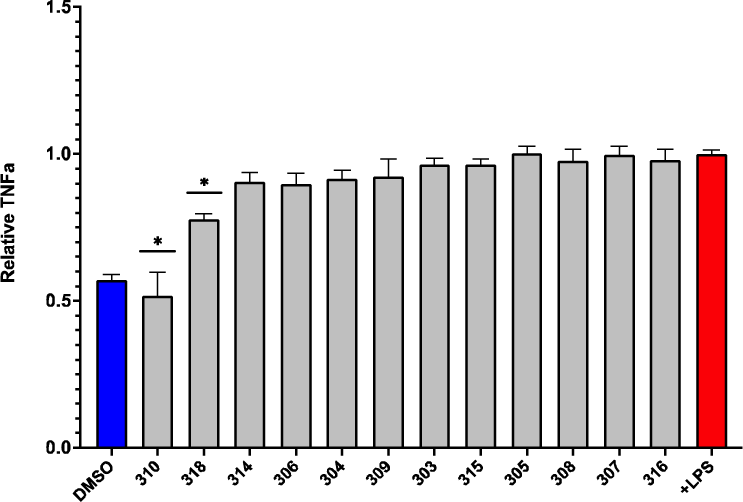
**Of the novel tricyclic congeners to FDA-approved tricyclic drugs which inhibit LPS-induced microglial TNFa secretion, only GM310 and #318 significantly reduced LPS-induced microglial TNFa (only compounds which were not toxic were included in the analysis).**

We then further assessed the protective effects of these novel tricyclic compounds to delay Abeta proteotoxicity in the *C. elegans* model of AD. As shown in **Figure 7**, five delayed delay proteotoxicity at Day 15 in the *C. elegans* model of AD, but only GM310 also inhibited TNF-a secretion.

**Figure 7.**
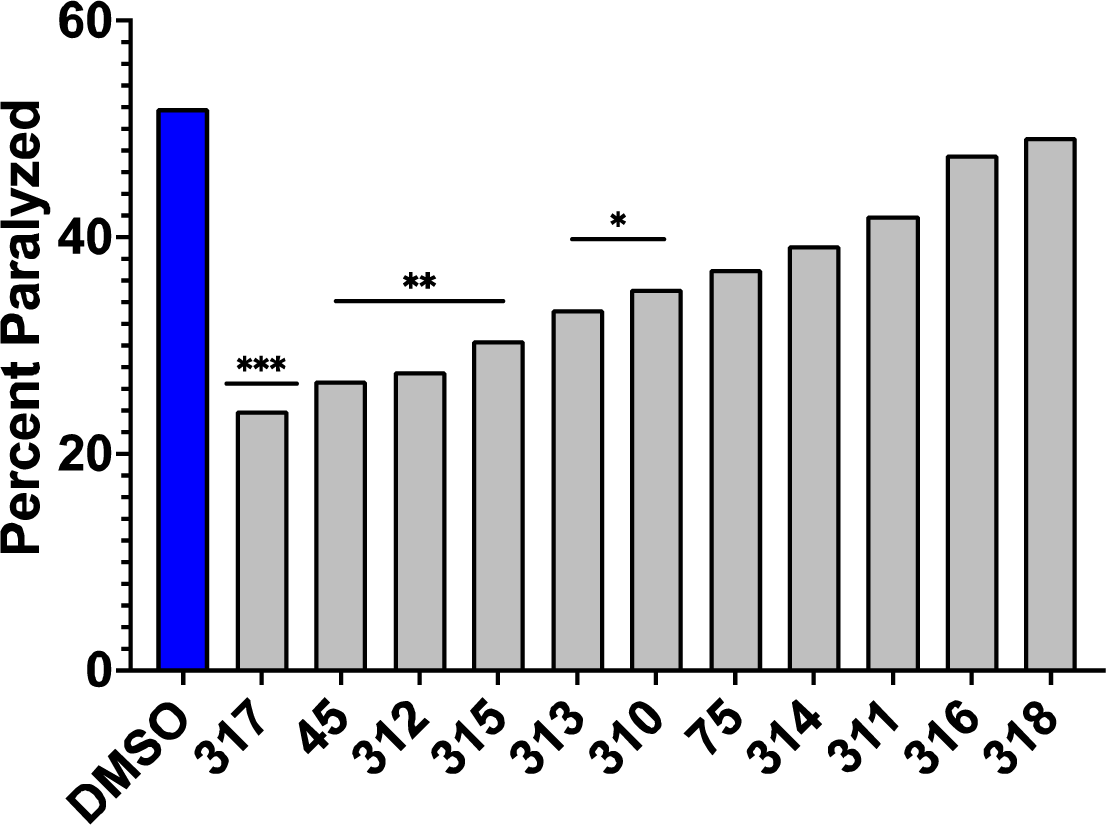
Of the novel tricyclic congeners, #317, 312, 315, 313, and 310 significantly inhibited Abeta proteotoxicity at day 15 in the C. elegans model of AD (*p<0.05,Chi-squared). Only GM310 inhibited both microglial TNFa secretion AND Abeta proteotoxicity.

We therefore carried out a more extensive dose-response analysis of the effects of GM310 over the entire lifespan of the CL2006 strain assessed every day until Day 24 of the adult phase. As shown in **Figure 8**, GM310 produced a robust dose-dependent delay in paralysis in this *C. elegans* model of AD, although at the highest concentration the compound was no longer protective. We have observed such behavior in most of the dose-response studies we have carried out in many different assays, and have interpreted such results as indicating some toxicity at the highest doses. This interpretation is consistent with the FDA requirement that dose-response studies be carried for a sufficiently wide range of doses to be able to calculate not only ED50 but also LD50.

**Figure 8.**
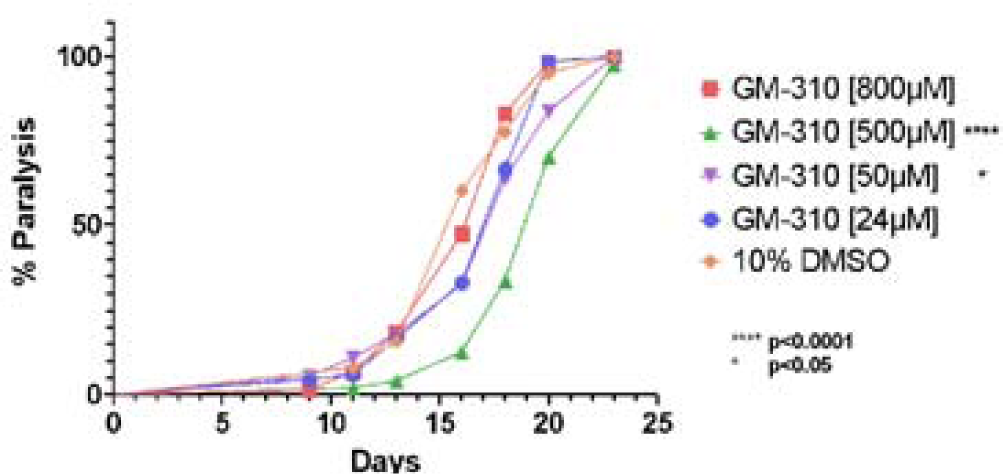
Novel tricyclic compound GM310 produces dose-dependent delay in proteotoxicity the *C. elegans* model of AD.

Since AD entails at least two forms of proteotoxicity (Abeta and Tau), and likely other proteins as well, we have focused on the functional impairments produced by transgenic expression of the human Abeta 1-42 peptide, rather than the levels of Abeta or Abeta aggregates. However it is still possible that the protective effects of the tricyclic compounds could be due to reduction of some aspect of Abeta which we did not measure. To address this issue, we assessed the protective effects of several tricyclic compounds in a completely different model of proteotoxicity produced by the disease-causing allele of the human Huntingtin protein, which we have used to extend our mechanistic studies of Abeta proteotoxicity [122]. As shown in **Figure 9**, several tricyclic compounds which inhibited BV2 cytokine secretion and protected in the *C. elegans* model for AD, were also protective in the *C. elegans* model for Huntingtin proteotoxicity. These studies indicate that the tricyclic compounds are generally protective against common pathophysiological processes that drive proteotoxicity. Although the tricyclic compounds, including GM310, protect against proteotoxicity and inhibit microglial secretion of TNFa, a concern with small molecules is that they may nevertheless produce other off-target toxic effects which have not been assessed but could preclude clinical application. To address this concern, we examined effects of GM310 on lifespan in the wild-type (N2) strain.

**Figure 9.**
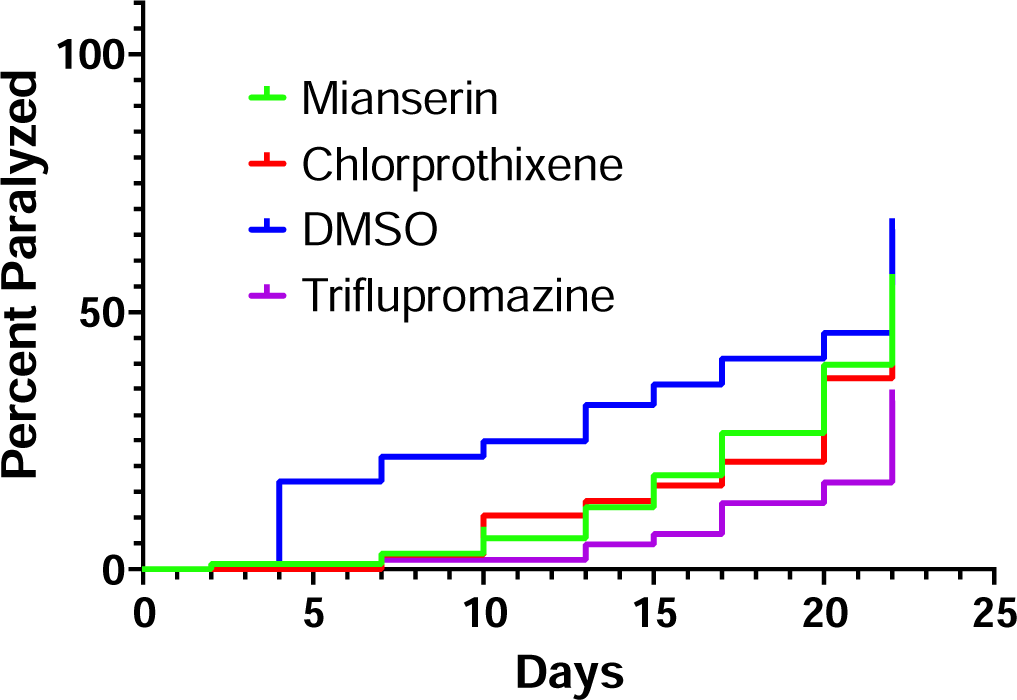
Tricyclic compounds which reduce microglial secretion of TNFa and delay proteotoxicity in a C. elegans model of AD also delay proteotoxicity in a C. elegans model of Huntington’s Disease.

As shown in **Figure 10**, GM310 significantly increased lifespan in wild-type *C. elegans*. This result indicates that even if this compound produces some off-target deleterious effect we simply have not yet detected, it produces protective effects that more than compensate, thus increasing lifespan.

**Figure 10.**
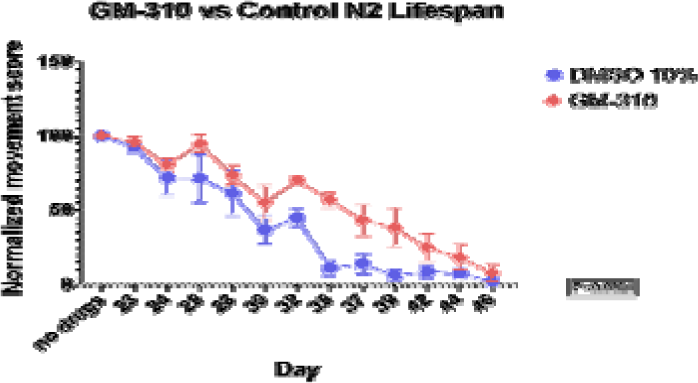
Novel tricyclic compound GM310 also increases lifespan in wild-type (N2) *C. elegans*.

To assess the potential efficacy of GM310 to treat dementia in humans, we assessed efficacy of the compound to ameliorate behavioral impairments, including memory, in a mouse model of intracerebral hemorrhage [123]. Although dementia due to AD is more common than vascular dementia, post-stroke cognitive impairment and dementia (PSCID) is a major source of morbidity and mo tality after stroke, including ICH [124]. As stated by Zhou et al., “These results confirm that AD and ICH may have common pathogenesis and share preventive treatment measures” [125]. Furthermore, assessing efficacy of interventions in mouse models of AD requires significant investments in time and funds, yet these models continue to be highly controversial [126]. In contrast mouse models of ICH are generally accepted as likely predictive of effects of interventions ICH in humans [123, 127, 128], and such studies require far less investments in time and funds than mouse models of AD, so more suitable to assess feasibility.

One of the major, yet often surprisingly overlooked, obstacles in developing drugs to treat neurological and psychiatric conditions, including dementia, is that “more than 98% of small molecule drugs and almost 00% of large molecule drugs are precluded from drug delivery to brain” [129, 130]. A major feature of the tricyclic compounds that have been approved to treat psychiatric conditions such as psychosis and depression is that in general and almost by definition these compounds cross the blood-brain barrier after oral administration [131]. We therefore assessed if compound GM310 would be concentrated in the brain after i.p. or oral delivery. As shown in **Figure 11**, GM310 is indeed highly concentrated in the brain after either i.p. or oral delivery. To assess the potential efficacy of GM310 to treat dementia, we assessed efficacy of the compound to ameliorate behavioral impairments, including memory, in a mouse model of vascular dementia caused by intracerebral hemorrhage (ICH) [123]. Vascular cognitive impairment (VCI) is the second most common type of dementia after Alzheimer’s disease (AD) [132]. Post-stroke cognitive impairment and dementia (PSCID) is a major source of morbidity and mortality after stroke, including ICH [124, 125, 133, 134]. Furthermore, vascular dementia and dementia due to AD appear to be driven by similar pathophysiological mechanisms, including a role for ApoE4 [135, 136] which entails inflammation, a mechanism common to these forms of dementia [137] and it has been concluded that interventions effective for one would likely be effective for the other [138, 139]. More specifically, ApoE4 promotes the secretion of microglial TNFa which in turn reduces neuronal viability, compared to other ApoE alleles [103]. This observation is particularly pertinent since ApoE is the major genetic risk factor for AD, and many studies have demonstrated that increased TNFa is a major contributor to the pathophysiology of both AD [29-40, 84, 85, 103] and stroke [104-107]. A different cytokine, IL-6, has also been implicated in driving the neuropathology of both AD and stroke [108], suggesting that dementia in both conditions might be driven by a general increase in inflammatory cytokines. NFkappaB, the key transcription factor which generally induces pro-inflammatory cytokines, has also been directly implicated in the pathophysiology of AD [109] and stroke [110], suggesting that an even wider range of cytokines might similarly promoted neurotoxicity in both conditions. We therefore assessed efficacy of GM310 to ameliorate impairments, including memory impairments, in a mouse model of ICH [123]. In this study ICH was produced by autologous infusion of whole blood into the mouse striatum, followed immediately by infusion of 10 nM GM310 into the same area. As shown in **Figure 12**, infusion of 10 nM GM310 completely prevented behavioral impairments and inflammation (increased TNFa) after ICH.

**Figure 11.**
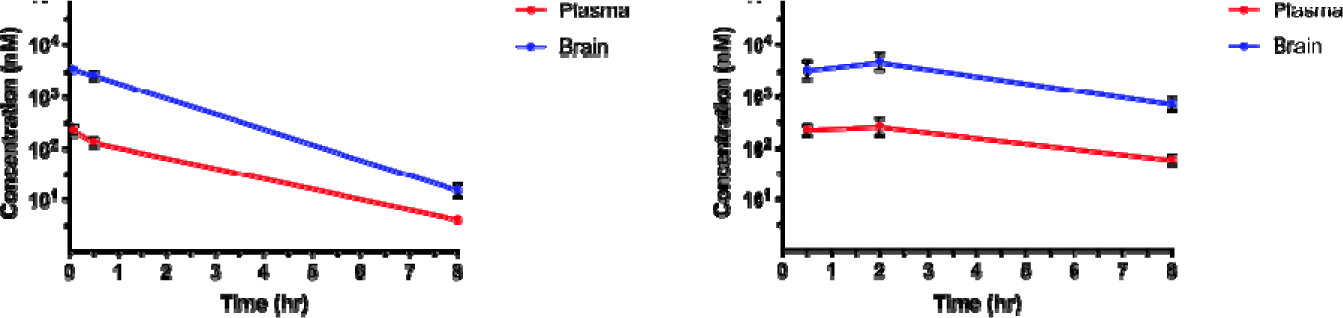
**Plasma and brain concentrations of GM310 over 8 hours following a single IV injection (A) at 1 mg/kg or oral administration (B) at 10 mg/kg in male C57Bl/6J mice. Plasma and brain (cortical) concentrations at each time-point represent the mean +/- SD from 3 mice. Arrow indicates effective dose of GM310 to block impairments in a mouse model of stroke (Fig. 10).**

**Figure 12.**
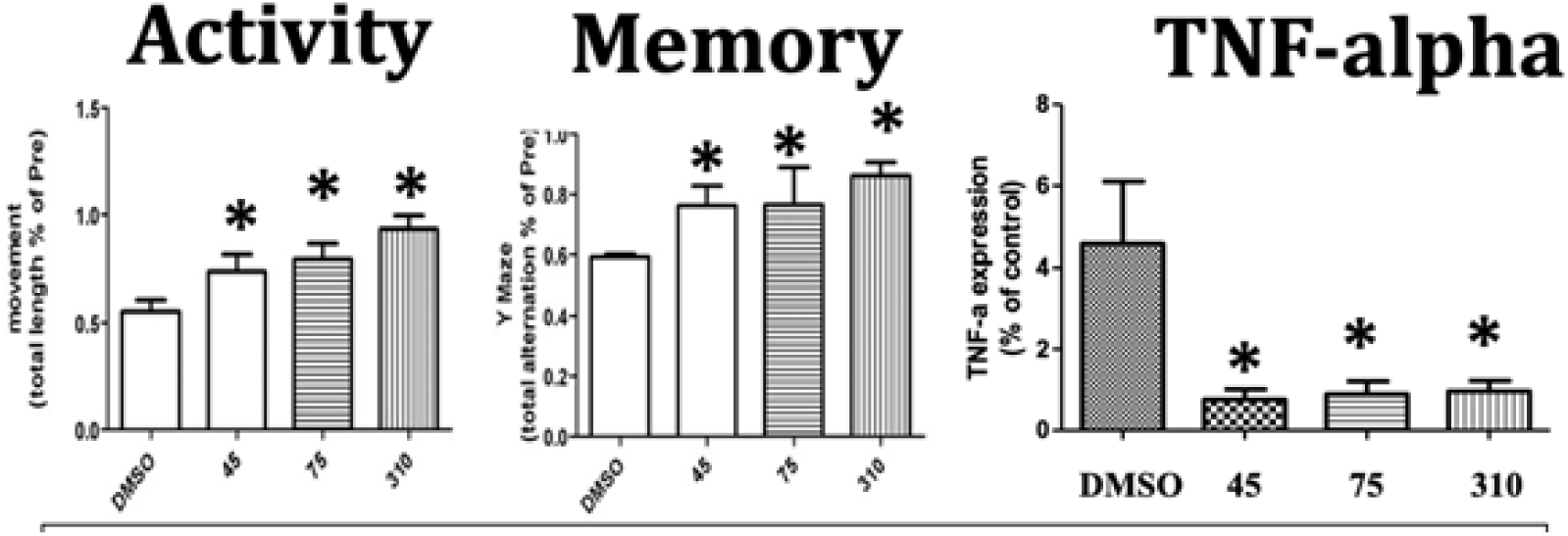
**Infusion of 10 GM310 into site of ICH completely protects against impairments in activity and memory and increased TNFa concentrations at the site of the ICH (*p<0.05).**

TNFa and inflammation in general have been implicated as driving pathophysiology of AD [29-40, 84, 85, 103] and stroke [104-107]. A different cytokine, IL-6, has also been implicated in driving the neuropathology of both AD [37] and stroke [108]. We therefore assessed the extent to which GM310 generally inhibits the microglial (BV2) secretion of pro-inflammatory cytokines, assessed by the Luminex Cytokine Array multiplex assay. As shown in **Figure 13**, GM310 inhibited microglial secretion of almost all proinflammatory cytokines in this assay, corroborating and extending previous studies focused on TNFa and IL-6, suggesting that GM310 might be clinically useful to treat a wide variety of conditions in which inflammation has been implicated, including dementia due to AD and vascular dementia (see above). The wide range of cytokines inhibited by GM310 is consistent with the hypothesis that GM310, like other tricyclic compounds, inhibits NFKappaB expression by directly inhibiting the transcription complex that stimulates NFKappaB expression [119].

**Figure 13.**
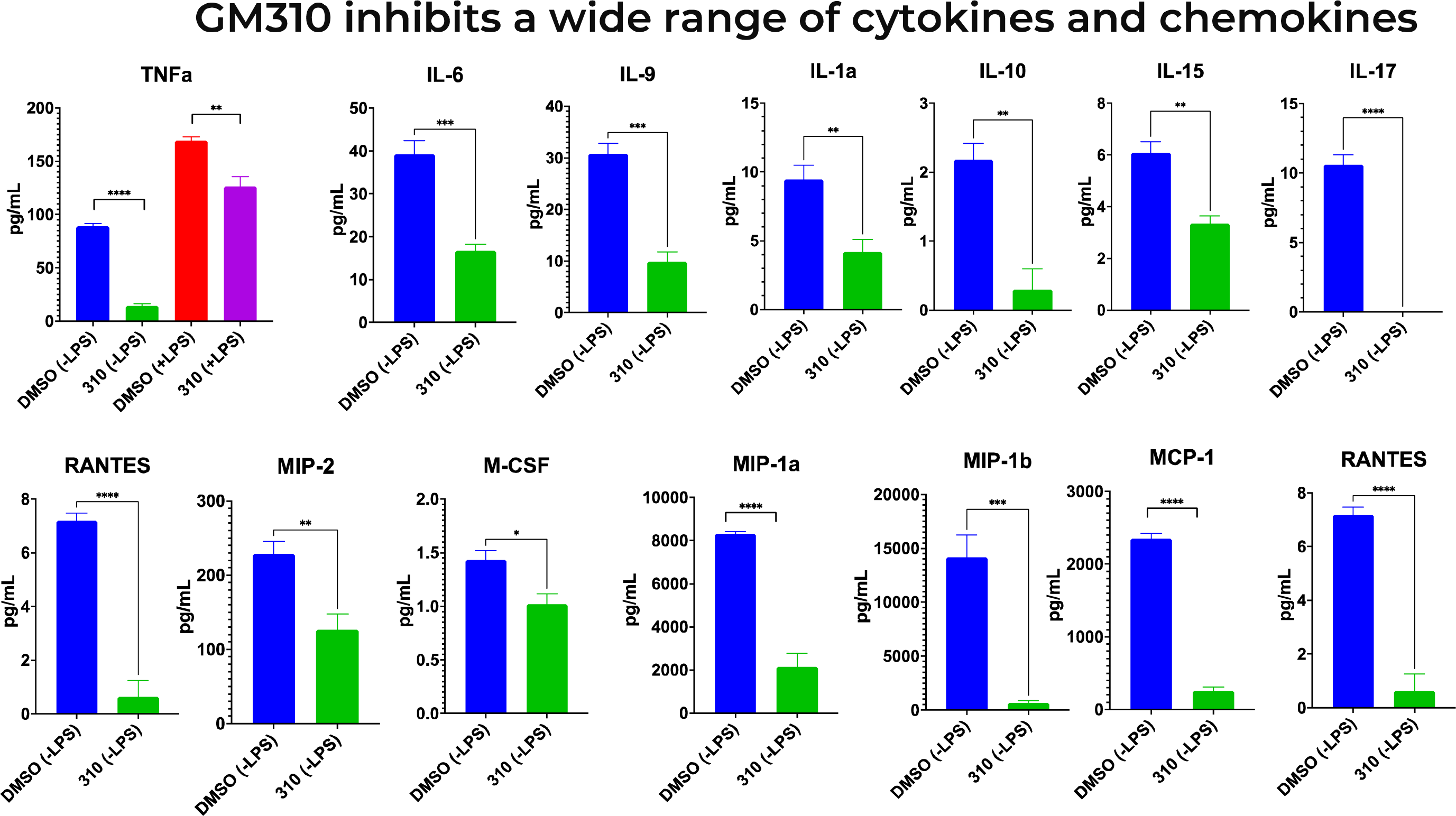
Novel tricyclic compound GM310 inhibits BV2 microglial secretion of cytokines. Supernatant was taken 48 hours after addition of 8uM drug and 24 hours of LPS stimulation using Luminex Cytokine Array multiplex assay. *** *P* < 0.0001, treated condition vs. LPS control.

Although it is now widely agreed that GWAS studies clearly implicate microglia gene expression as driving major aspects of the pathophysiology of AD [40], and as described above the evidence is overwhelming that enhanced microglial inflammation similarly drives major aspects of the pathophysiology of many forms of dementia [29-40, 84, 85, 103] and stroke [104-107], it is plausible that impaired microglial repair mechanisms, especially phagocytosis, could also contribute to these impairments [140]. Thus it is possible that if a compound reduces microglial inflammation but also reduces phagocytosis, the net effect could be that these processes in effect cancel each other out, or even worse, if impaired phagocytosis is quantitatively more important than enhanced inflammation, such a compound could conceivably worsen impairments in dementia. It is therefore critical to assess the effects candidate compounds which inhibit microglial inflammation on phagocytosis. We therefore assessed the effect of all the compounds which inhibited both Abeta proteotoxicity AND microglial cytokine secretion on microglial phagocytosis (10μg/ml pHrodo-red labeled zymosan (Thermo Fisher #P35364), 1:100 pHrodo-red labeled early apoptotic Jurkat (EAJ) cells, and 1:100 red fluorescent carboxylate-modified polystyrene latex beads (Millipore Sigma #L3030, 0.9 μm). The large majority of compounds which inhibit both microglial inflammation AND Abeta proteotoxity had no effect on phagocytosis (fluorescence normalized to MTT); thus at least these compounds are unlikely to worsen dementia symptoms, and presumably, due to favorable protective effects, are likely to ameliorate symptoms (**Fig. 14**). However, 2 of the tricyclic compounds, #228 and #246, did reduce phagocytosis, so would not be good candidates to develop to treat dementia (**Fig. 14**). Remarkably, the only tricyclic compound which inhibits microglia inflammation and also *increases* microglial phagocytosis is GM310 (**Fig. 14**), which further supports that this novel compound is the most plausible lead so far to treat dementia.

**Figure 14.**
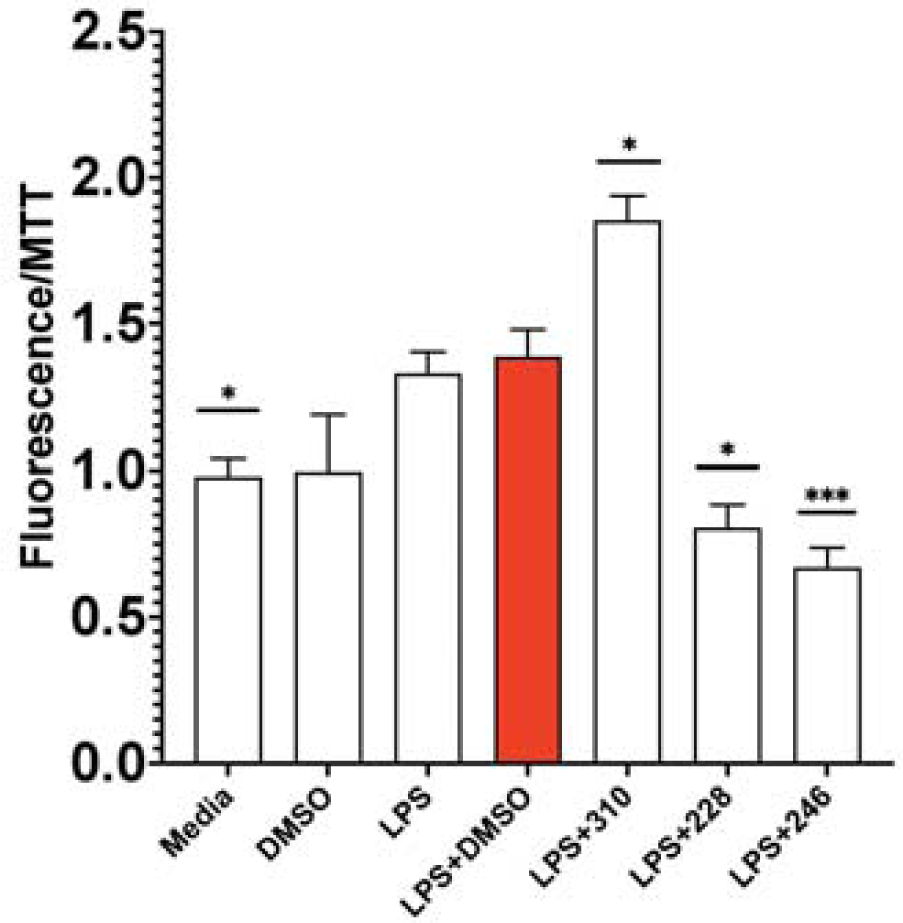
GM-310 is only compound tested which decreased inflammation and increased phagocytosis in microglia.

To assess potential efficacy of GM310 and its congeners to treat dementia in human patients, we assessed if these compounds would also inhibit LPS-induced TNFa secretion in human primary monocytes (**Figure 15**) and human PBMCs (**Figure 16**). As shown in **Figure 15**, almost all of the test compounds significantly inhibited TNFa from human monocytes. Similar results were obtained using PBMCs isolated directly from blood of patient volunteers (**Figure 16**). Of particular interest, GM310 was the most effective compound to reduce TNFa in human PBMCs.

**Figure 15.**
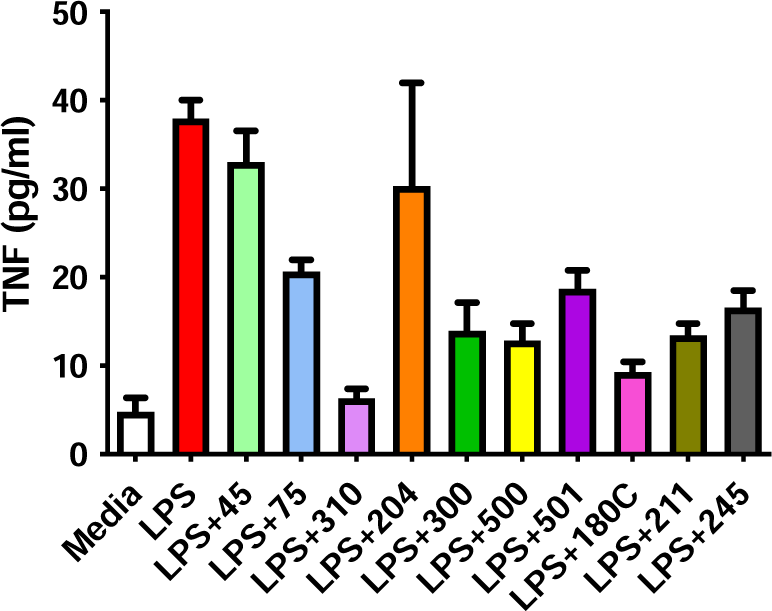
GM310 and congeners inhibit LPS-induced secretion of TNFa in human primary monocytes (effects of compounds 75, 310, 300, 500, 501, 180C, 211, and 245 all p<0.01).

**Figure 16.**
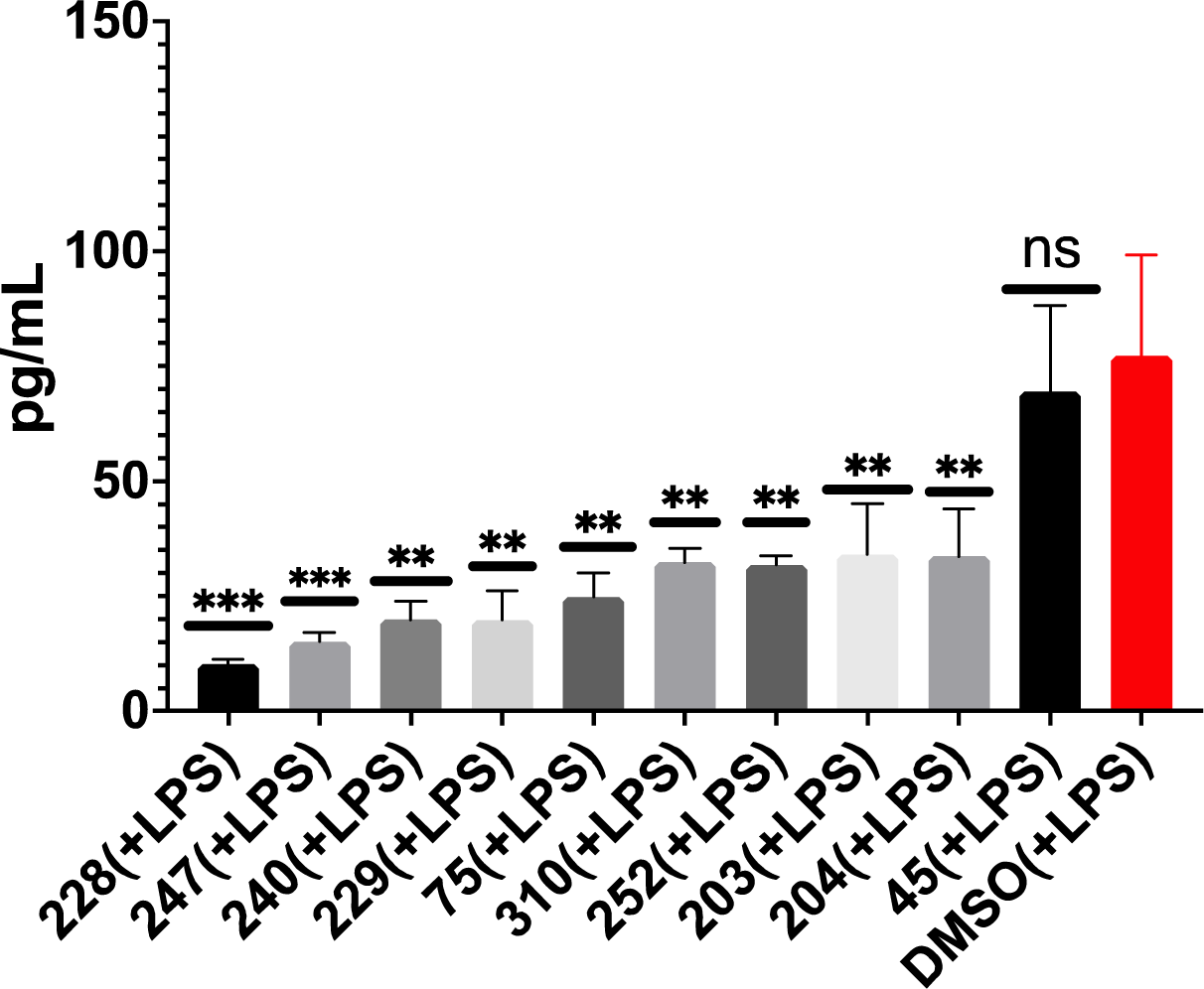
GM-310 and congeners inhibit LPS-induced secretion of TNFa in human PBMCs.

To clarify mechanisms mediating these broad protective effects of GM310 and its FDA-approved tricyclic congeners, we assessed effects of these congeners on gene expression in human neuronal cell lines using the CMAP/L1000 database [141]. Although of course GM310 was not in the database, since it is novel, the effects of 14 tricyclic congeners of GM310, FDA-approved tricyclic compounds zapproved by the FDA to treat either psychosis or depression, on gene expression in 2 neuronal cell lines (NEU and NPC) were available in the database. (Ideally we would have carried out the analysis in a human microglial cell line, but such a line does not exist in this database, and even if it did, it is doubtful that effects of psychoactive drugs on microglial gene expression would have been made a high priority).

Using a composite Z-score to estimate the net effects of these tricyclic compounds on gene expression, we observed that the only genes inhibited by all 14/14 tricyclic compounds as indicated by the CMAP database, were the transcriptional co-inhibitor related genes CtBP2 and CtBP1. CtBP has long been of interest to us because its transcriptional co-inhibitor activity is mediated at least in part by inhibition of Cbp activity [142], which as indicated above, mediates protective effects of dietary restriction on lifespan and Abeta proteotoxicity [143], is allosterically activated by glucose metabolism (specifically via NADH) which we have argued is the metabolite most responsible for deleterious effects of glucose metabolism during aging and diabetes [96]. In turn NADH allosterically activates CtBP, while promotes inflammation by enhancing transcriptional activity of NFKappaB on inflammation by allosteric inhibits the transcriptional co-activator CBP, whose induction we showed mediates the protective effects of dietary restriction on lifespan and proteotoxicity in the CL2006 C. elegans model of Abeta proteotoxicity [52], subsequently corroborated in mice [52] and flies [144]. We also showed that hypothalamic expression of CtBP inversely correlates with lifespan [52] consistent with reports that neuronal CtBP shortens lifespan [145], the opposite of CBP [146]. As a co-transcriptional inhibitor, CtBP also recruits HDACs to the chromatin to inhibit transcription, and as described above, we [146] and others have demonstrated that HDAC inhibition is highly protective in a wide range of models relevant to dementia [61, 147-150]. It is therefore of great interest that CtBP2 gene expression is significantly elevated in almost every brain region examined in patients with AD, compared to controls, according to the AGORA database (Sugis et al, 2019). the most inhibited gene, significantly inhibited by 10/14 FDA-approved tricyclic congeners of GM310, was cysteine-rich protein 1 (CSRP1), which is conversely highly induced in all brain cell types, including microglia, in brains of Alzheimer’s patients, as indicated by the scREAD database [151], as well as in a mouse model of AD (Beckmann et al., 2020) and has been nominated as a potential target for AD by the agora.ampadportal.org (Ming et al., 2021). Of particular interest, the reversal of age-related human cardiovascular impairments by FOX03, the human analog of daf-16, appears to be mediated by inhibition of CSRP1 [152].

To further assess potential mechanisms mediating anti-inflammatory effects of GM310 and other tricyclic compounds in microglia, we assessed gene expression in BV2 microglia 24 hours +/LPS, +/- 8 uM GM310, using RNAseq. Remarkably, the gene expression profile produced by LPS alone was, as expected, characteristic of increased glycolysis and expression of several cytokines, including TNF-a, whereas GM310 produced a gene expression profile highly similar to that which we had shown is produced by dietary restriction in mouse hypothalamus, including induction of Creb-Binding Protein (CBP, p<10^-5^), Catalase (Cat, p<10^-14^), and Snf2-related CREBBP activator protein (Srcap, p<10^-3)^, with inhibition of genes which oppose Cbp activity such as HDACs, and a robust shift in gene expression indicating a shift toward metabolism of alternative substrates relative to glycolysis (e.g., increased CPT1) (**Figure 17**).

To assess molecular mechanisms mediating the anti-inflammatory effects of GM310 in microglia, we assessed effects of GM310 on gene expression in LPS-stimulated BV2 microglia by RNAseq. Remarkably, effects of GM310 on microglial gene expression were highly similar to those produced by dietary restriction in the hypothalamus [143]. Specifically, GM310 induced Cbp, which have demonstrated to be required for the protective effects of dietary restriction on lifespan and Abeta proteotoxicity [143], and the effects of dietary restriction on metabolic gene expression indicating reduction in glucose metabolism and induction of metabolism of alternative substrates.

To functionally assess if this change in gene profile also functionally produced a reduction in glycolysis, we assessed these functions in BV2 microglia using flux analysis (Seahorse Fluxometer) 24 hours +/- LPS, +/- tricyclic compounds including GM310. As demonstrated in **Figure 18**, LPS induced glycolysis, as expected, and GM310, inhibited glycolysis with or without LPS. Further analysis demonstrated that almost all tricyclic congeners examined prevented the increase in LPS-induced BV2 microglia glycolytic capacity (not shown).

**Figure 18.**
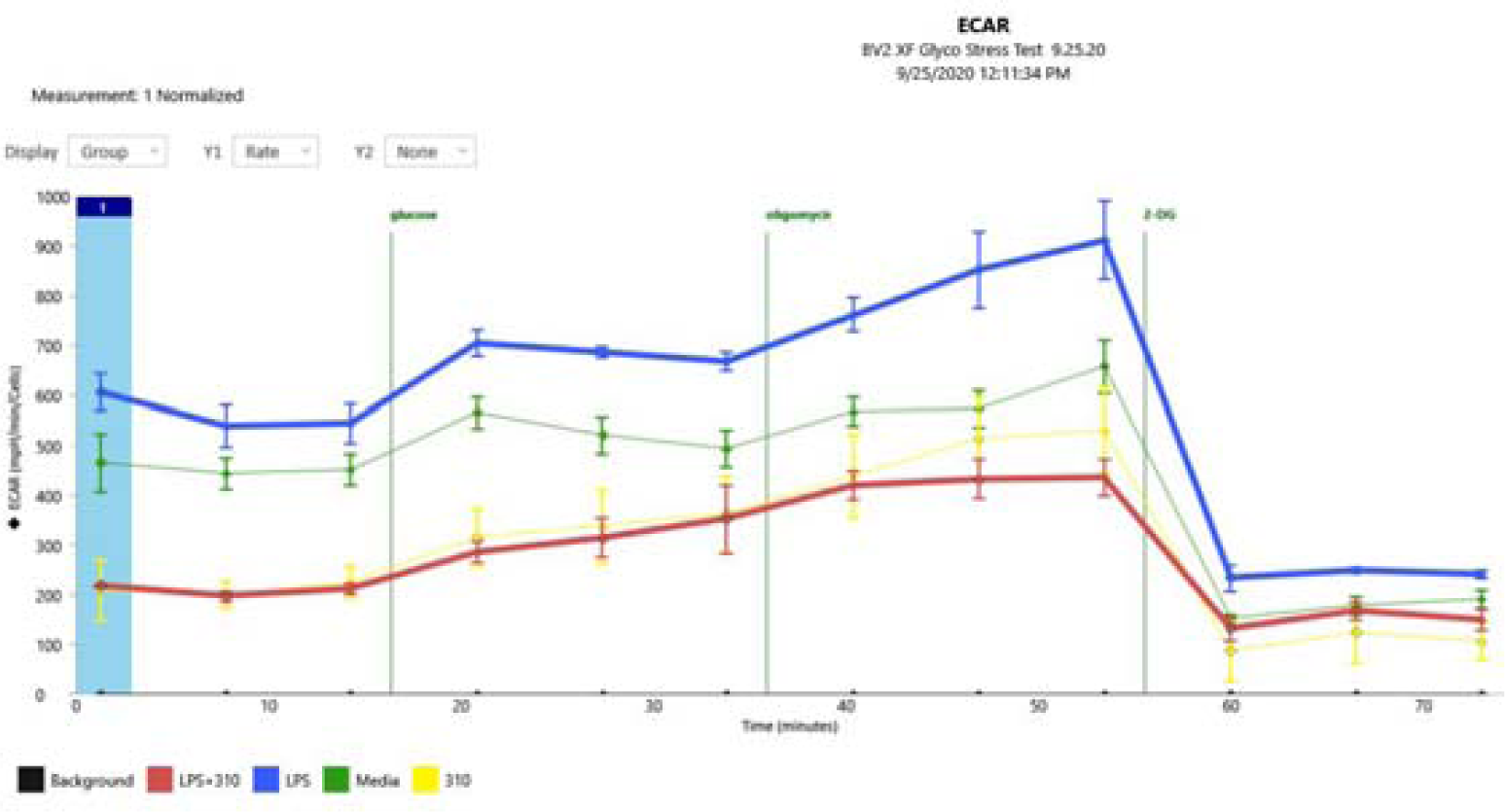
**LPS induced glycolysis in BV2 cells, and compound GM310 reduces glycolysis with or without LPS.**

To more fully assess the role of glycolysis in driving Abeta proteotoxicity, we assessed the effect of reducing glycolysis by the hexokinase inhibitor 2-deoxglucose (2-DG) on paralysis in the CL2006 *C. elegans* model. As shown in **Figure 19**, 2-DG significantly delayed Abeta proteotoxicity and increased lifespan in wild-type C. elegans.

**Figure 19.**
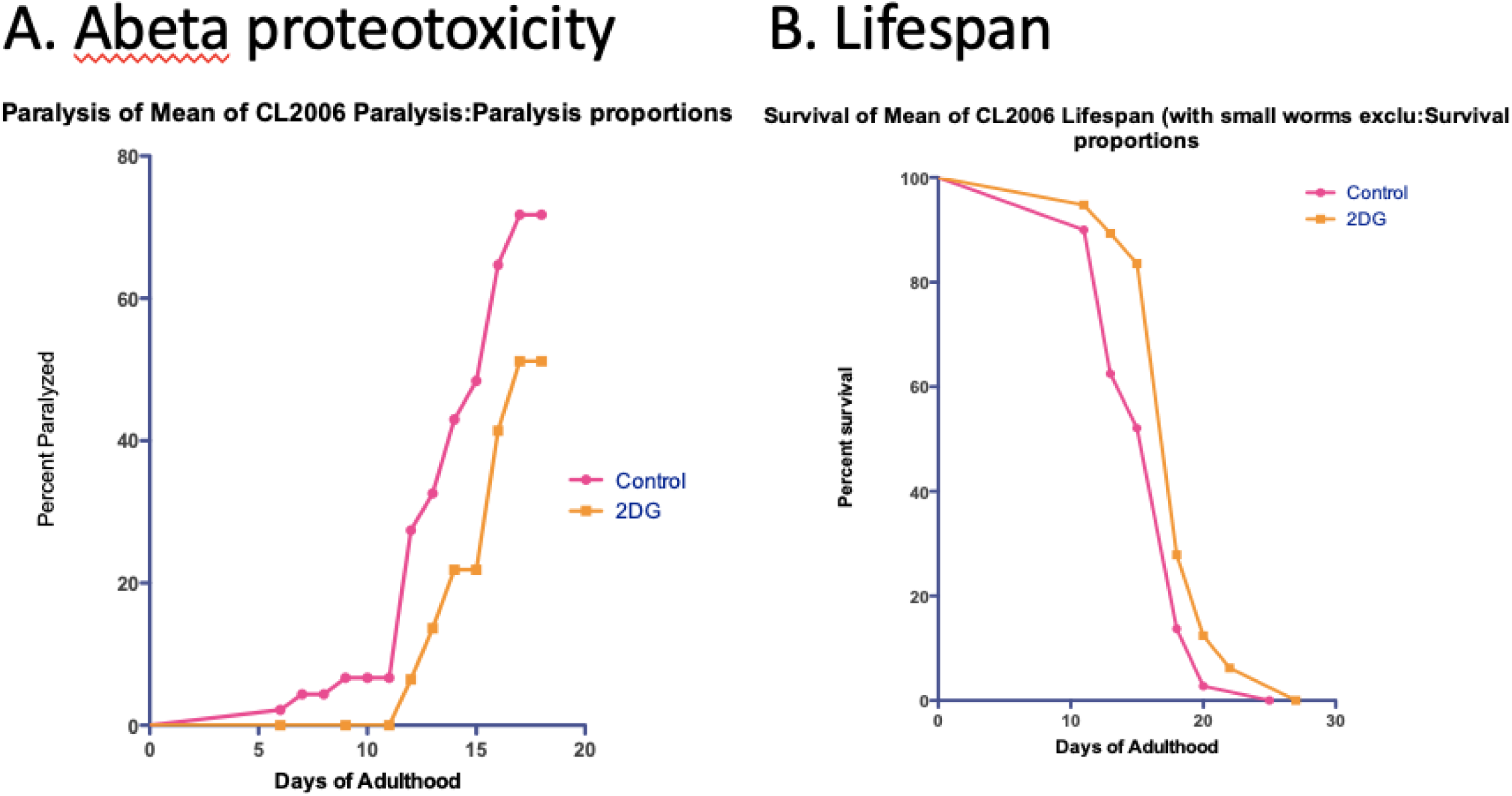
**Glycolytic inhibitor 2-deoxyglucose (2-DG) significantly delays Abeta proteotoxicity in the CL2007 strain and increases lifespan in wild-type C. elegans.**

## DISCUSSION

The purpose of the studies reported here was to discover small bioavailable molecules which could mimic the protective effects of dietary restriction to generally delay age-related diseases [153], discovery of which it was hypothesized, would lead to novel elucidation of mechanisms mediating these protective effects, leading development of even more effective treatments. Successful prosecution of such a program requires appropriate choices of phenotype that reflect the general goal (e.g., general delay of age-related diseases in humans) while allowing high-throughput screening. A far more common strategy to develop such therapies has been to focus on specific targets hypothesized to be a critical driver of a disease (e.g., Abeta peptide for Alzheimer’s Disease), and develop compounds acting on that target. Although this is the standard approach taken, the history of drug discovery indicates that first-in-class effective drugs (penicillin, anti-psychotics, etc.) have almost never been discovered, by such a target-based approach, almost by definition definition (successful drugs lead to mechanism, rarely vice-versa). Instead, such drugs have generally been discovered on the basis of phenotype, often noted by alert clinicians (e.g., anti-psychotic drugs).

An alternative plausible approach to discover drugs that are broadly effective against age-related diseases has been to carry out high-throughput screens to discover compounds [54] or genes whose inhibition increase lifespan [154-156]. The model organism *C. elegans* constitutes a particularly effective platform for carry ng out such high-throughput screens, so the extent to which mechanisms and interventions which delay age-related disease phenotypes in *C. elegans* apply to humans is particularly important to validate the use of this model. To date results have been encouraging-the discovery of the role of insulin-like signaling in determining lifespan in C. elegans, as produced by inactivating mutations in the *daf-2* gene [157, 158] has been shown to be equally important in determining lifespan in flies [159], mice [160, 161], and probably even humans [162-166]. Furthermore, RNAi inhibition of *daf-2* delays development of disease-related impairments in the CL2006 *C. elegans* model of AD used in the present study [113] even when instituted a relatively advanced age at which this construct no longer extended lifespan [167], an observation particularly relevant to potential therapies. Dietary restriction similarly delays AD disease-related impairments in the *C. elegans* CL2006 model, an effect mimicked by the HDAC inhibitor butyrate and dependent on the histone acetyltransferase Creb Binding Protein (Cbp) [52]. Dietary restriction is similarly protective in mouse models of AD [168-172], possibly mediated by similar molecular mechanisms mediating protective effects of *daf-2* inhibition [173].

Consistent with the role of Cbp in mediating protective effects of both dietary restriction and inactivation of *daf-2* in lifespan [52, 63, 146, 174-176] and against AD disease-related phenotypes in *C. elegans* [52, 122], Cbp is also protective in mouse models of AD [177]. However, we have observed that molecular and pharmaceutical interventions which increase lifespan in *C. elegans* rarely delay disease-related impairments as in the CL2006 model for AD (Gonzalez et. al, unpublished). Therefore, in the present study we focused on phenotypes, proteotoxicity and inflammation, implicated in driving age-related diseases, rather than lifespan.

The results of these studies strongly support that the phenotypes chosen for the screening protocol used are highly effective to discover compounds likely to be protective in human AD. The 3 most protective compounds, of 2560 screened in the CL2006 strain, were each individually or in a class of drugs already demonstrated to be highly protective in a wide range of models for neurodegenerative diseases, especially AD: phenylbutyrate [56-62] [100], methicillin [66-69], and quetiapine [70, 101, 102]. Phenylbutyrate combined with taurursodiol (a bile acid with unclear mechanism of action) significantly improves symptoms in patients with ALS [178, 179] and is now approved by the FDA to treat this condition. The same combination is currently in clinical trials to treat AD, with promising results so far on surrogate measures https://www.amylyx.com/pipeline. It is also of some interest that, based on the CMAP database [141], phenylbutyrate alone reversed the expression of 36% of the genes differentially expressed in microglia in AD [151]. Quetiapine combined with valproate, like phenylbutyrate an HDAC inhibitor, also improves symptoms in patients with AD [180].

That these or related compounds would protective in AD was even more strongly supported by their robust inhibition of microglial TNF-a (**Figures 3-5, 13**), and human myeloid cells (**Figures 15,16**) strongly implicated by both genetic and interventional studies in driving neurological diseases, including in patients with AD [26, 27, 29–40, 46, 84, 85, 181]. Conversely, impaired microglial phagocytosis has also been implicated in driving AD [140], and GM310 at least enhances microglial phagocytosis (**Figure 14**), in addition to broadly inhibiting pro-inflammatory cytokines (**Figure 13**). Furthermore, as indicated above, a purely bioinformatic analysis had also indicated that quetiapine should be effective in treating AD [71]. Taken together these data strongly support that the novel compound GM310, which also crosses the blood-brain barrier at human therapeutic oral doses (**Figure 11**), and completely protects in a mouse model of stroke (**Figure 12**), is likely to be protective in human AD as well as stroke and other neurodegenerative diseases. Although potentially off-target deleterious iatrogenic effects are always a concern with any therapeutic intervention, the observation that GM310 increases lifespan suggests that if there are toxic effects (Figure 10), they are more than compensated by systemic protective effects.

An equally important outcome of the present studies is the elucidation of the likely mechanism by which these protective effects are exerted. Although protective and anti-inflammatory effects of phenybutyrate, quetiapine, and the other compounds studied herein could be mediated by a wide range of targets, the observation that GM310 induces changes in gene expression highly similar to that produced by dietary restriction (**Table I**), and is protective against models of diseases as disparate as AD (**Figure 8**) and stroke (**Figure 12**), and increases lifespan (**Figure 10**), and structural similar compounds were protective in a C. elegans model of Huntington’s Disease **(Figure 9)** suggests that GM310, and plausibly other compounds, exert their protective effects through mechanisms similar to those which mediate dietary restriction. Although many mechanisms have been hypothesized to mediate the broadly protective effects of dietary restriction to delay age-related impairments have proposed, a fair reflection of the field is that these mechanisms include inhibition of mTOR and insulin-like activity (thus overlapping with protective effects of inhibiting daf-2 and equivalent activities in mammals and other species), and increase of sirtuin activity [182]. While many studies have addressed mechanisms by which these activities influence age-related impairments and lifespan, there is no consensus, and for that matter little speculation, about how these disparate mechanisms mediate protective effects of dietary restriction.

However, based on a wide range of data, we have proposed that protective effects of dietary restriction are mediated by reduction in glucose metabolism [13, 52, 94, 96, 183-188], and all mechanisms generally proposed for dietary restriction plausibly act through this final pathway. This hypothesis was originally based on the pattern of gene expression produced by dietary restriction or hypoglycemia in the hypothalamus, which strongly indicated that dietary restriction and hypoglycemia reduce glycolysis and induce metabolism of alternate substrates such as lipids (beta oxidation, as indicated by induction of Cpt1a) and ketones [185], observations which were later corroborated by much more extensive validation with DNA microarrays followed by quantitative PCR [189]. Thus these corroborating studies also demonstrated that dietary restriction inhibited expression of genes which promote glycolysis (e.g., phosphofructokinase, and glucose transporter 2), or transport of glycolytic products into the mitochondria (malate dehydrogenase), while inducing genes which inhibit glycolysis (e.g., pyruvate dehydrogenase kinase) or reverse glycolysis (pyruvate decarboxylase), and promote metabolism of alternative substrates (e.g., Cpt1a, fatty acid transporter, patin-like phospholipase) [189]. Of particular interest, dietary restriction induced the expression of hypothalamic FOXO3 [189], the mammalian ortholog of daf-16, whose induction is required for the protective effects of inactivating *daf-2*, as well as for glycolytic inhibition to increase lifespan [190].

We observed that dietary restriction produces the same metabolic switch, away from glycolysis and toward utilization of alternative substrates, in whole-body *C. elegans* [52]. In the same study we observed that several transcription factors in the hypothalamus were also either induced or inhibited by dietary restriction or hypoglycemia, and the expression of one transcription, Creb-binding protein (Cbp) positively correlated with lifespan over 5 strains of mice [52]. The C. elegans ortholog of Cbp was also induced by dietary restriction, and inhibition of Cbp by RNAi prevented the metabolic switch away from glycolysis and toward alternative substrates and prevented the protective effects of several forms of dietary restriction as well as the daf-2 mutation to increase lifespan and delay impairments in the CL2006 model of AD [52]. Since Cbp is a histone acetyltransferase (HAT), we further demonstrated that inhibition of HDAC activity by sodium butyrate, which opposes HAT activity, mimicked the protective effects of dietary restriction to increase lifespan and delay impairments in the CL2006 model of AD, which was also blocked by inhibition of Cbp [52]. Although not reported in that paper, we also observed that the transcription co-inhibitor, CtBP, which also opposes Cbp activity [142], was inhibited by dietary restriction, and expression was inversely correlated with lifespan (Poplawski et al., unpublished). It was subsequently reported that viral-mediated increased expression of Cbp increased BDNF and reduced learning and memory impairments in a mouse model of AD [177], another example of results from *C. elegans* predicting disease mechanisms in vertebrates.

Since GM310 produces highly similar molecular responses as produced by dietary restriction (**Table 1**), the most plausible mechanism mediating the protective effects of GM310 is the same as that mediating protective effects of dietary restriction, specifically reduction in glucose metabolism. We directly corroborated this prediction (**Figure 18**), and further confirmed that direct inhibition of glucose metabolism by 2-deoxyglucose (2- DG) did indeed mimic the protective effects of GM310 to delay impairments in the CL2006 strain, and increased lifespan in the wild-type strain (**Figure 19**). We had previously demonstrated that direct inhibition of glucose metabolism by 2-DG also delayed impairments in a *C. elegans* model of Huntington’s Disease [122]. Consistent with these results, 2-deoxyglucose was protective in mouse models of Parkinson’s Disease [191] and stroke [192]. Similarly, caloric restriction in yeast is produced by simply reducing glucose concentrations, which increases extends life-span in yeast, associated with activation of sirtuin activity, which is activated by NAD+ which in turn is reduced by glycolysis [193-195]. Reducing dietary glucose also increased maximum lifespan in *Drosophila* [196]. Consistent with these observations, reducing glucose metabolism directly by the hexokinase inhibitor 2-deoxyglucose increases replicative lifespan in human fibroblasts [197]. Based on these and other studies, it has been proposed that small molecules which mimic the protective effects of dietary restriction can be classified into two roughly equal classes, as “upstream”, which effectively reduce glucose metabolism, and “downstream”, which may mediate protective effects of reducing glucose metabolism [198]. For example, D-glucosamine increases lifespan in both *C. elegans* and mice [199], plausibly by inhibiting glycolysis [200]. Methionine increases lifespan associated with reduced blood glucose [201]. Similarly we demonstrated that compounds which protect against glucose toxicity increase lifespan [174], and the ketone 3- hydroxybutyrate (3-OHB, which appears to mediated the reversal of diabetic nephropathy by elevating this ketone, also protects against glucose toxicity [202]; we have now also demonstrated that the ketogenic diet reverses age-related kidney failure (Isoda et al., unpublished). 3-OHB also increases lifespan in C. elegans associated with reduced glucose toxicity [203]. Indeed, possibly the earliest lead compound used to develop small molecules to mimic protective effects of dietary restriction was 2-DG [204]. Similarly, it has been proposed that inhibition of glycolysis is a particularly promising approach to developing small molecules which mimic the protective effects of dietary restriction [205, 206], possibly even in primates including humans [207]. Further important support for this hypothesis is that rapamycin, the compound which produces the most robust increase of lifespan so far [208-212], increases replicative lifespan in apparently by blocking the increased glycolysis which occurs in this process [213]. Similarly, recent studies have demonstrated that the amino acid taurine decreases with age in humans, and taurine supplementation produces robust increased lifespan in mice and healthspan in monkeys [214] and is protective against cognitive impairments in aging humans, and in mouse models of AD and Parkinson’s disease [215], and reduces inflammation, plausibly by reducing glycolysis [216].

**Table 1.**
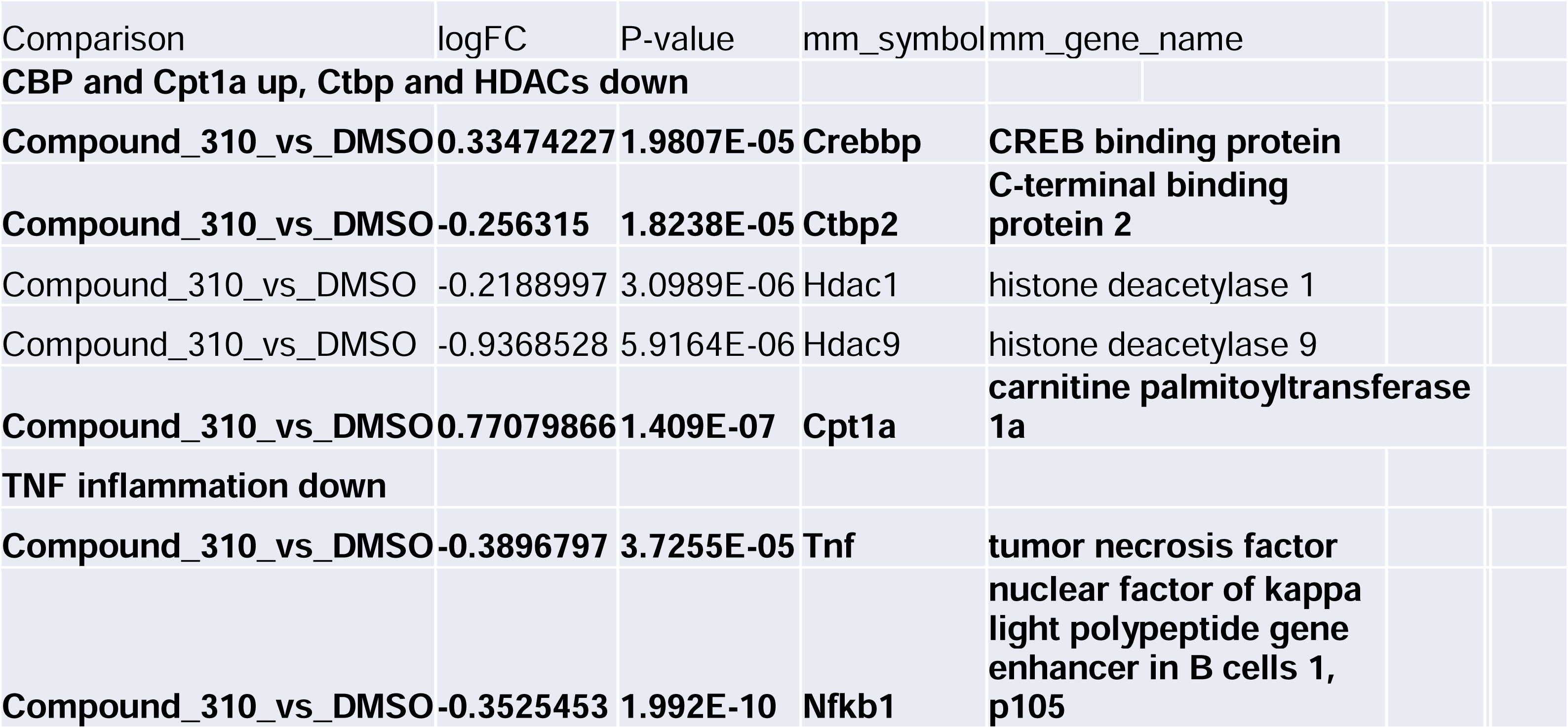
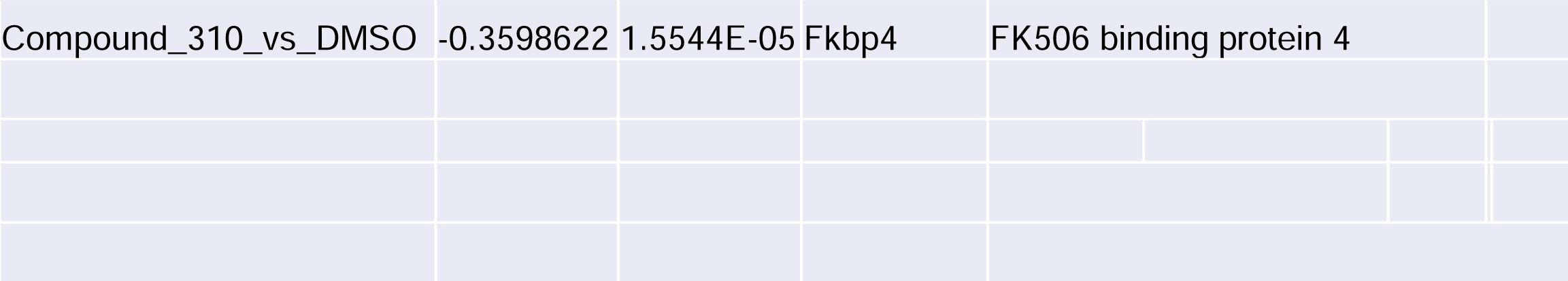
Effects of GM-310 on microglial gene expression is similar to effects of dietary restriction on hypothalamic gene expression: i duction of Cbp, inhibition of opposing HDAC gene expression and CtBP2 expression, and a gene profile promoting metabolic pathway genes, suc as beta oxidation relative to glycolysis. Consistent with the metabolic profile, TNF-alpha and pro-inflammatory gene expression are inhibited. Sign of logFC indicates induction vs. inhibition.

Conversely, it is well-known that whole-body glucose metabolism varies even over the 24-hour cycle, as measured by indirect calorimetry, with whole-body glucose metabolism decreasing after each meal and rapidly increasing after each meal as plasma glucose increases [217], so molecular responses to dietary restriction and hypoglycemia in the hypothalamus simply reflect the metabolic state of the whole animal. That glucose metabolism drives age-related impairments is strongly supported by clinical and animal studies demonstrating that increased glucose drives toxic processes in diabetes associated with increased glucose metabolism {Keen, 1994 #5232;Van den Enden, 1995 #6860; #5233;Greene, 1999 #5067;Pop-Busui, 2010 #8099}, and in particular shortens lifespan and inducing glycolysis by other interventions also increases oxidative stress and is toxic [218]. Dietary restriction is also highly protective to reduce the development of cancer [219-229], most likely by reducing glycolysis on which cancer depends (referred to as the Warburg effect) [12]. Dietary restriction also robustly inhibits inflammation [6, 7, 169, 171, 230-232], also most likely by inhibition increased glycolysis, which is required for inflammation [10, 97]. In fact it has been proposed that protective effects of dietary restriction during aging are mediated by inhibition of inflammation [230] through a metabolic mechanism [7]. Of particular interest to the present studies, diabetes is a major risk factor for AD [233-237] as well as stroke [138, 238, 239], further supporting the GM310 protects in animal models of both conditions by decreasing glucose metabolism.

A major reason for that glucose metabolism is more toxic than that of alternative metabolites is that that most of the energy to produce ATP from glycolysis is derived from the transfer electrons from NADH to ETC Complex I, then transfer to ETC Complexes III, IV, and V, which are the main sources of reactive oxygen species, but much less so from ETC complex II, [240-245]. This basic fact about mitochondrial function explains why inactivating mutations in complex II produce profound sensitivity to oxidative damage as well as reduced life span [246, 247]. Genome-wide screening studies have demonstrated that genes coding for mitochondrial functions constitute possibly the most conspicuous single class of ‘senescence assurance genes’, ablation of which increases life span [156, 248]. Almost all of these life-span-limiting mitochondrial genes code for proteins in mitochondrial complexes I, III, IV or V [156, 248] [155] [154]. For example, of 23 genes discovered in an exhaustive genome wide screen whose inhibition increased life span [154], 12 were genes coding for proteins in mitochondrial (ETC) complexes I, III, IV or V, and one gene coded for a key enzyme in glycolysis, glucose-6-phosphate isomerase. An independent screen from another laboratory obtained very similar results, though discovering a different glycolytic enzyme whose inhibition increases life span [154]. Strikingly absent from these screens were genes for proteins in mitochondrial complex II [[154, 155]. Indeed, classic genetic screens had already identified that mutations causing impairments in complex II reduce life span [249]. Thus proteins in mitochondrial complexes I, III, IV and V and at least some glycolytic enzymes function to limit life span, whereas genes for proteins in mitochondrial complex II function to increase life span, consistent with the preferential toxicity of glucose metabolism vs. that of other metabolites. Furthermore, dietary restriction diverts glucose from glycolysis, which drives oxidative stress, to the pentose pathway, for example by inducing transketolase [250]. The pentose pathway produces NADPH, the main source of antioxidant reducing power in the cell [251, 252], as opposed to the production of NADH, ultimately the main source of oxidative stress in the cell. In addition to increasing lipid metabolism, e.g., by inducing Cpt1a, dietary restriction also increasing protein turnover to allow metabolism of amino acids, both of which thus function to reduce lipids and proteins which may have incurred damaging modification due to oxidation. Thus by reducing glucose metabolism, dietary restriction (and by inference GM310) reduces oxidative stress, increasing anti-oxidant defenses, and reduces lipids and proteins which may have been damaged by reactive oxygen species. Furthermore by converting NAD+ to NADH, glycolysis reduces the levels of NAD+, which thereby reduces the activity of the protective effects of sirtuins, also strongly implicated in the protective effects of dietary restriction [194, 195, 253]. Similarly, the production of NADH activates CtBP, inhibition of which increases lifespan probably by enhancing lipid metabolism [254] by antagonizing Cbp [142]. These phenomena thus explain why reduction of glucose metabolism, and activation of alternative metabolic pathways, by dietary restriction is protective during aging.

Although many toxic processes have been proposed to mediate impairments with age, these mechanisms generally only address acute toxicity without addressing why impairments increase with age, other perhaps than there are cumulative effects of such toxic events (i.e., that they are a consequence of the second law of thermodynamics). In contrast, glucose metabolism as a driver of age-related impairments could also explain cumulative effects because glucose leads to persistently-induced glycolysis in a phenomenon referred to as “metabolic memory” [255, 256] [256-264]. We have specifically demonstrated that exposure to elevated glucose produces persistently increased glucose metabolism, which is not reversed after restoration of normal glucose, but is prevented by the 3-OHB and Ppar agonists [256], which also reduce glucose metabolism [265] (indeed the physiological function of Ppar-alpha is to produce the shift away from glycolysis toward lipid metabolism during fasting [266]). This phenomenon may thus explain why inflammation increases with age [2], in turn leading to the many age-related conditions driven by increased inflammation, including neurodegenerative diseases like Alzheimer’s (see above). The induction of glucose metabolism by glucose, associated with reduced metabolism of alternative substrates, is in accordance with one of the main tenets of biochemistry, which is that substrates induce their own metabolism and inhibit alternative metabolic pathways, as canonically demonstrated by the lac operon [267]. Interestingly, the induction of the lac operon also exhibits persistent effects, as we [256], and others have demonstrated for glucose [255, 256] [256-264]. Of further interest, the induction of the lac operon by lactose exhibits stochastic bistable behavior [268], and similar behavior in driving persistent effects of glucose could also explain why stochastic processes appear to play a major role in determining lifespan in genetically identical individuals under virtually identical environments [96], as observed in *C. elegans* [269], and even in human twin studies [270] [271].

While the cumulative toxic effects of glucose metabolism during aging could be cell-autonomous, it is possible that the the age-related increase in glucose metabolism that occurs in at least some tissues (e.g., myeloid inflammatory and cancer cells) could be mediated by a small group of nutrient-sensing neurons which regulate whole-body glucose metabolism [13]. This hypothesis is supported by the observation that protective effects of the induction of Cbp by dietary restriction are mediated by nutrient-sensitive GABA neurons [146] and that the protective effects of dietary restriction in *C. elegans* are mediated by two just two nutrient-sensing neurons [272]. Furthermore, nutrient-sensing neurons in the hypothalamus, which sense both glucose and the hormone leptin which reflects long-term energy balance as reflected by adipose stores, regulate whole-body glucose production and metabolism [273-278].

The remarkably broad protective effects of tricyclic compounds indicated by the present studies as well as previous studies might seem surprising. However, it should be emphasized that the therapeutic efficacy of this class of compounds has largely been discovered through serendipity. Methylene blue, the archetypal example of this class of compounds, is considered to be the first synthetic compound widely used for medicinal purposes, and whose medicinal efficacy was originally assessed because of its selective staining of specific organelles and even pathogens, although it became clear early on that while methylene blue did indeed exhibit protective medicinal effects, these effects are entirely independent of its staining properties, and in fact the staining properties have been eliminated to facilitate use of its congeners. The availability of many methylene blue congeners facilitated the discovery of the first anti-histamine, benadryl, and the use of the side effect of benadryl as a soporific to calm agitated schizophrenic patients led to the serendipitous discovery that the congener of benadryl, chlorpromazine, reduces negative symptoms of schizophrenia, leading to the pharmacopeia of tricyclic congeners of chlorpromazine now used to treat psychosis and depression. Indeed, further studies in this vein have demonstrated that methylene blue itself exhibits properties likely to be protective in AD and even more broadly other age-related diseases [279-283]. Thus the independent discovery of the protective effects of tricyclic compounds in these studies further supports the value of the phenotypic screen protocols developed for this purpose.

An interesting further implication of these observations is that the antipsychotic and antipsychotic effects of tricyclic compounds could be mediated by their anti-inflammatory effects, since these psychiatric conditions disorders are associated with increased inflammatory responses [86, 284]. These observations could also explain why AD increases the risk these psychiatric conditions [72] [75, 78].

## Author Contributions

RL developed the approach, carried out the screening in *C. elegans* and cells, analyzed data, and wrote the manuscript. JV screened BV2 cells, BTZ screened *C. elegans,* LS, K-HR, AS, GM, NP and YM carried out bioinformatics; YY and CK carried out mouse stroke studies; HK, SV, and JJ synthesized novel compounds; IR-l carried out studies in human PBMCs; and CM developed the approach and wrote the manuscript.

## Methods

### Microsource Spectrum library

The Microsource Spectrum library of 2560 compounds, includes all of the compounds in the US and International Drug Collections, in addition to the Microsource Natural Product and Discover libraries. The collection provides biologically active and structurally diverse compounds. The Microsource Spectrum library of compounds was provided as 10mM stock solutions in DMSO and distributed in a plate by the Mount Sinai high-throughput core facility. All drugs were diluted to and tested in our screens at the concentration of 8μM.

### Strains and genetics

All strains were maintained at 20 C as previously described. *Caenorhabditis elegans* strains used were: Bristol strain (N2), AD muscle model (CL2006). All strains were obtained from the CGC

### *C. elegans* – maintenance and preparation of L4/adult stage

The *C. elegans* CL2006 Alzheimer’s muscle model strain was maintained at 20°C on nematode growth medium (NGM) agar seeded with *Escherichia coli* strain OP50. Chemical synchronization was performed to eliminate variation caused by age differences. Worms at the L4/adult stage were plated in the experimental plates containing liquid medium, *Escherichia coli* strain OP50, FUDR and the drugs to be screened.

### *C. elegans* - Paralysis assay

Primary screening experiment: Paralysis was assessed in liquid medium in 384-well plate liquid cultures maintained at 20°C. Age synchronized *C. elegans* were plated as L4/young adults (6-12 worms per well) in plates containing drugs dissolved in DMSO, S Complete Medium, *E.Coli* OP50. To prevent self-fertilization, 5- fluoro-2’-deoxyuridine (FUDR) was added to the liquid medium at a final concentration of 120μM. A small shock derived from a 4.5V battery is applied to each well for <1sec to stimulate movement of the worms. Each well was recorded for 90 seconds after worm stimulation. Videos were analyzed to assess paralysis, defined as the absence of a full sinusoidal movement of a worm after stimulation.

Secondary assays: Setup, maintenance, analysis are similar to primary screen. Differences include: 96-well plates were used for all secondary assays (10-15 worms per well). Drugs were added a few hours after worms were placed in the liquid medium plates.

For the initial screen, videos were taken and paralysis was assessed on Day 12 or 15 (with control displaying >50% worms paralyzed). Subsequent paralysis analysis was conducted by analyzing worm movement every other day from Day 5 in addition to Day 15 analysis.

### *C. elegans* -Lifespan assay

Preparation for the lifespan assay was similar to the paralysis assay and L4/ young adult N2 wildtype worms were placed in plates containing liquid medium and FUDR, drugs were added a few hours after plating the worms. The fraction of alive animals was based on a movement score assessed with the WormMicrotracker (wMictoTracker) 5 days a week.

### *C. elegans* Microtracker

The MicroTracker measures overall locomotor activity and viability of worms using a laser beam reader. We assessed worm movement 30 minutes 5 days/week until control had a movement score less than 50 mouvements.

### Statistical analysis

Analysis was completed using the Prism software using the log-rank test for comparisons and P-value comparisons for paralysis assays and 2-way Anova for lifespan analysis.

### Cell culture studies/Cytokine ELISA

LPS (Invitrogen 00-4976-93), PBS and ELISA kits (TNFa and IL-6) were purchased from Invitrogen. MTT (M2128) was ordered through Sigma-Aldrich. All screened compounds were ordered through the National Cancer Institute Developmental Therapeutics Program.

### BV2 murine microglia

BV2 were cultured in Dulbecco’s Modified Eagle Medium supplemented with 5% FBS and Penicillin Streptomycin and all experiments were done with cells below passage 30. To induce an inflammatory response, microglia were exposed to LPS (1.25- 10ug/mL). Following stimulation or drug treatments, culture media was collected for detection of secreted cytokines with Invitrogen ELISA kits and cell viability was determined using MTT assay. LPS was diluted in 10% DMSO to prepare stock solution to be added to complete culture media. DMSO concentrations were no more than 0.1%.

BV2 microglia were plated 5000 cells per well in a 96 well plate and incubated at 37°C overnight. Media was replaced with fresh media supplemented with stimulants (amyloid beta or LPS) and incubated at 37°C. Compounds were added directly to the existing media at a final concentration of 8uM and incubated at 37°C. Supernatant was collected for cytokine analysis with ELISA and cell viability determined using MTT assay.

### TNFa and IL-6

were measured in cell culture supernatant using ELISA (Invitrogen). Absorbance was read using a Thermo Scientific MultiSkan Go at 450nm.

Cell viability was determined using MTT assay. Culture media supplemented with MTT (0.5mg/mL) was added to each well and incubated for 1.5 hours at 37°C. Media was aspirated and replaced with 100uL of 4% HCl in Isopropanol to lyse cells and dissolve MTT. Absorbance was measured using a Thermo Scientific MultiSkan Go at 560nm and 620nm. Optical densities were normalized to the PBS treated wells to determine cell viability. All statistical analyses and graphs were generated using GraphPad Prism software. Statistical analysis for drug and stimulation conditions included a one-way analysis of variance with Dunnett’s Post Hoc test to determine individual differences among conditions.

### Glycolytic flux

was measured using the Seahorse XF96 Analyzer (Seahorse Bioscience) as we have previously described [256], at the Mt. Sinai Mitochondrial Analysis Facility.

## Acknowledgements

Our special thanks to the Mount Sinai Human Immune Monitoring Center, Live Imaging and Bioenergetics Facility, and MSIP (especially Louise Lammers, Dov Shamir, and Elaine Lu), and Joan Horvath for developing system to monitor *C. elegans* activity.

We particularly thank Dr. Rudy Tanzi for scientific advice and Drs. Jack Rowe, John Morrison, Patrick Hof, Peter Rapp, James Roberts, Don Pfaff, Caleb Finch, Steve Kleopoulos, Eric Nestler, and Paul Kenny for long-standing support without which these studies could not have been done.

This work was supported by a pilot grant and an award from MSIP, and by NIH R01AG062303.

